# Signaling induced biophysical disruption of repressed chromatin domains drives immune cell fate

**DOI:** 10.64898/2025.12.08.692984

**Authors:** Alexia Martínez de Paz, Christopher R. Chin, Mythili Ketavarapu, Yuchen Sun, Matthew R. Marunde, Jeremy T. Chang, Laiba Khan, John Cohen, Rui Yang, Michael J. Bale, Andrew W. Daman, Vigneshwari E. Kumar, Chenyang Jiang, Dughan J. Ahimovic, Mark Owyong, Arjun Ravishankar, Wilfred Wong, Rochelle Shih, Bria Graham, Catherine E. Smith, Ioannis Karagianidis, Leandro Cerchietti, Christopher R. Flowers, Michael R. Green, Christopher E. Mason, Joseph C. Sun, Alejandro Martin-Trujillo, Rachel E. Niec, Yicheng Long, Michael-Christopher Keogh, Shixin Liu, Wendy Béguelin, Christina S. Leslie, Ari M. Melnick, Steven Z. Josefowicz

## Abstract

Cell fate transitions require signal-induced chromatin derepression, yet mechanisms governing transitions from repressed to active chromatin states are poorly understood. We discover, at fate-defining genes across immune cell types, a signal-induced histone code, and describe domains of H3 serine 28 phosphorylation (H3S28ph) spanning architectural features, often coincident with repressive H3 lysine 27 trimethylation (H3K27me3). Employing biophysical, single cell, and functional approaches to study signal-induced cell differentiation in the immune system, we uncover epigenomic transitions and cell fate choices precipitated by histone phosphorylation (H3ph). Mechanistically, H3ph overrides Polycomb Repressive Complex 2 (PRC2) chromatin repression, biophysically disrupts polynucleosome compaction, and promotes loss of H3K27me3, while increasing activating H3K27 acetylation and H3K36 dimethylation to drive domain interactivity and stabilize transcription. We demonstrate the activity of H3ph in several cell fate transitions and illuminate biophysical mechanisms enabling rapid signal-activated chromatin derepression, processes with general relevance for cellular differentiation and activation.

## Introduction

Cell fate transitions rely on the activation of signaling cascades, transcription factors (TFs), and epigenetic mechanisms. These initiate specific transcriptional programs and are often characterized by rapid kinetics. Many studies have examined TFs activated by signaling pathways, focusing on their regulation, target genes, and interactions with transcriptional machinery^1,2^, with TFs widely considered the terminal nodes in signaling pathway activity. Less understood is how signals are directly transduced to chromatin and how the resulting chromatin states contribute to transcriptional activation. It is widely accepted that TFs associate with and guide chromatin modifying co-activators to convert chromatin to a transcriptionally permissive state. However, we largely lack mechanistic insights into how stimulation induced genes are rapidly derepressed, particularly in physiologic contexts of cell stimulation, such as immune cell differentiation or activation. Phosphorylation of histone proteins couples signal transduction to rapid transcriptional responses^3–5^. In response to external cues, histone H3 phosphorylation is rapidly (within minutes) deposited at inducible target genes^5^. Histone phosphorylation has been proposed to coordinate with adjacent lysine acetylation and (as in the original histone code hypothesis^6^) act as a “switch” to antagonize neighboring lysine methylation^7^. These models are supported by the correlation of histone phosphorylation with transcriptional activation, its interference with chromatin repressor complex binding, and its recruitment or stimulation of transcriptional co-activators^5,8–17^. Despite this, the identification of relevant chromatin states in primary cells and the biophysical mechanisms and dynamics of signal-activated chromatin transitions have remained elusive. Further, we lack an understanding of the prevalence and physiologic contexts of such chromatin states and the transcription programs and cellular and developmental fates they might control.

Immune cell activation, such as rapid activation of macrophages, T cells, B cells, and natural killer (NK) cells, serves as a highly relevant model for dissecting epigenetic mechanisms driving signaling-dependent chromatin state transitions. In addition, the germinal center (GC) reaction represents an ideal model for investigating these processes because the varied quality of signaling inputs directs B cells to undergo antibody affinity maturation, positive selection, and differentiation into memory B cells or plasma cells^18^. GC B cell phenotype is maintained through a tightly controlled repressive chromatin landscape where the deposition of H3K27me3 by PRC2 is critical^19,20^. GC B cell positive selection and fate transitions require a rapid signal-induced derepression of PRC2 target genes, including several bivalent loci which display both H3K27me3 and the activation-associated H3K4me3^21^. Despite the critical function of developmentally regulated H3K27me3 and bivalent chromatin in GC B cells, embryonic stem cells^22–24^, hematopoietic stem cell renewal and immune cell development^25–27^, and multiple other biological contexts, the chromatin mechanisms driving rapid de-repression are unknown.

Previously, we described the stimulation-induced phosphorylation of histone H3 serine 28 (H3S28ph) and histone H3.3 serine 31(H3.3S31ph) downstream of the MAPK and NF-kB signaling pathways, which promote rapid and robust transcription of macrophage inflammatory genes^9,10^. Here we define general principles of stimulation-responsive chromatin in diverse immune cell subsets and identify conserved molecular features of rapid domain-level chromatin state transitions. Further, we define the biophysical mechanisms of H3ph-mediated chromatin domain activation, and the generalized application of these pathways in instructing immune cell development and activation.

## Results

### Discovery of signal-activated chromatin states across immune cell types

Previously, we demonstrated that inflammatory gene induction in macrophages relies on the phosphorylation of H3S28 and H3.3S31^9,10^. These marks display distinct deposition patterns that we propose act synergistically in chromatin activation, with H3.3S31ph marking gene bodies and 3’ ends, consistent with its co-transcriptional nature, and H3S28ph mostly occurring within intergenic regions as well as enhancers and promoters. To better understand these marks’ deposition patterns, associated chromatin states, and to guide mechanistic models for their function in chromatin activation, we generated a series of epigenomics datasets in resting and stimulated macrophages and naïve B cells. These datasets included HiC for chromatin architecture context and ChIP, CUT&RUN, and/or CUT&Tag for H3S28ph, H3.3S31ph, H3K27ac, H3K27me3, H3K4me3, H3K4me1, H3K36me3, and H3.3 for chromatin state analysis employing chromHMM^28^ to integrate multiple epigenomic maps and annotate the genome based on the combinatorial patterns of chromatin marks. First, we observed broad stimulation-dependent deposition of H3S28ph spanning expansive intergenic chromatin domains and a clear delineation of topologically associating domains (TAD) in both macrophages and naïve B cells (**Figures 1A**, **S1A**). Stimulation induced H3S28ph and H3.3S31ph in both naïve B cells and macrophages shared general features but localized to cell-type and stimulation-specific response genes. In comparison with other common histone posttranslational modifications (PTMs), H3S28ph domains were comparable in breadth to those of repressive H3K27me3 (**Figure 1B**), highlighting the broad distribution of this modification, a feature common in repressive PTMs but unusual for an active chromatin associated PTM. H3.3S31ph peaks exhibited an intermediate peak size similar to co-transcriptional H3K36me3 (with which it colocalizes^10^) and broader than the promoter restricted H3K4me3. A primary goal of generating these datasets and chromHMM analysis (**Figure 1C**) was for unbiased discovery of chromatin states uniquely featured in stimulated cell states. We detected well-established chromatin states that define active enhancers, transcribed regions or H3K27me3 repressed chromatin, among others (**Figure S1B**). Inclusion of stimulated cell states in our analysis enabled the discovery of three previously undescribed chromatin states enriched with H3ph: induced transcribed regions (defined by dual H3.3S31ph and H3K36me3), H3S28ph induced regions and H3K27me3 repressed chromatin that gains H3S28ph after stimulation, which we term “transitional chromatin” (**Figure 1D**, **S1B**). Importantly, this unbiased chromatin state analysis shows that: (1) induced states are characterized by the dynamic deposition of H3ph, with the other PTMs being stable in the time frame analyzed; and (2) looking at the induced states in both cell types, H3S28ph emission parameters are highest where there is overlap with H3K27me3. The transitional chromatin state also displays H3.3 and H3K4me1 signal, suggesting overlap with poised enhancers. These regions often contain genes known to be rapidly induced upon activation of macrophages (*Socs1*, *Txnrd1*) and naïve B cells (*Myc*, *Cd86*) (**Figure 1E**, **S1C**). To understand if this newly discovered transitional chromatin state is a generalizable feature of the cellular response to stimuli, we profiled the deposition patterns of these marks in response to stimulation in naïve CD4 T cells and natural killer (NK) cells. As in macrophages and naïve B cells, we observe domain-level, stimulation induced co-enrichment of H3K27me3 and H3S28ph overlapping key master regulatory genes in both cell types. These include genes encoding key transcription factors in CD4 T cells (T-bet, NFATc1) (**Figure 1F**) and simulation-induced NK genes such as *miR155hg*, *Il2ra* and *Il15ra* (**Figure 1G**). Overall, our analysis discovered stimulation induced chromatin states and identified a prominent and common “transitional chromatin” state featuring repressive H2K27me3 and activating H3S28ph co-localized across large architectural chromatin domains containing critical activating and fate determining genes across multiple immune cell types.

**Figure 1.**
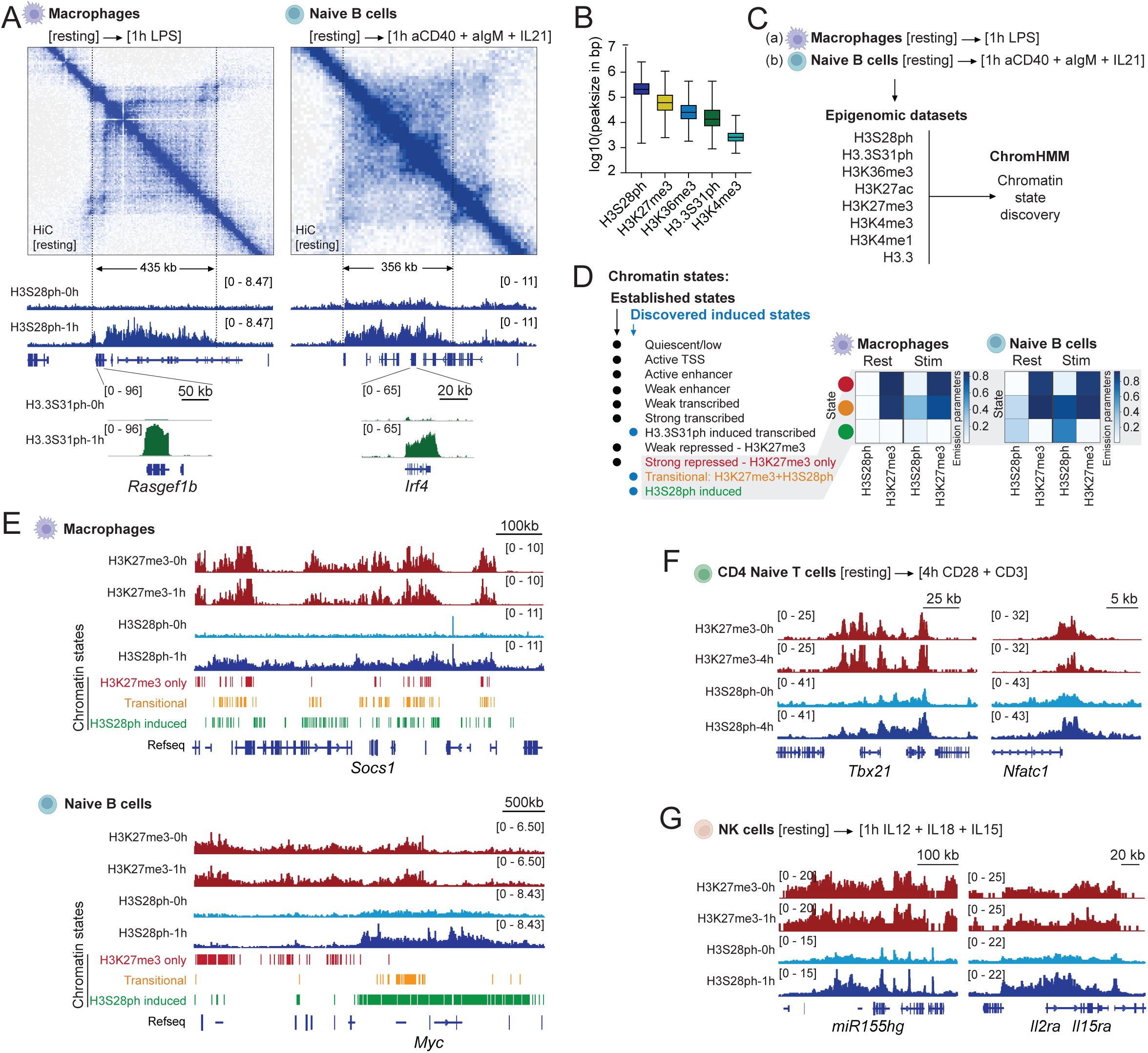
Discovery of signal-activated chromatin states across immune cell types. **(A)** Top: HiC contact matrix around *Rasgef1b* in resting bone marrow derived macrophages and *Irf4* in naïve B cells. Bottom: example Integrated Genomic Viewer (IGV) tracks from H3S28ph and H3.3S31ph ChIP experiments in resting and stimulated macrophages and naïve B cells. **(B)** Peak length distributions for different histone posttranslational modifications in naïve B cells. **(C)** Schematic of datasets generated for ChromHMM. **(D)** Chromatin states discovered by ChromHMM in resting and stimulated macrophages and naïve B cells. **(E)** IGV tracks for H3K27me3 and H3S28ph and highlighted chromatin states detected around *Socs1* in resting and stimulated macrophages and *Myc* in resting and stimulated naïve B cells. **(F)** Example IGV tracks for H3K27me3 and H3S28ph in resting and stimulated naïve CD4 T cells and **(G)** natural killer (NK) cells.

### Chromatin integrates signaling quality through the deposition of histone phosphorylation at induced genes upon germinal center B cell positive selection

Deeper analysis of H3S28ph and H3.3S31ph (H3ph) in naïve B cells showed that the top loci gaining H3ph include genes essential for germinal center (GC) formation and GC B cell positive selection, including *Myc*, *Irf4* and *miR155hg* (**Figures S2A, B; Table S1**). Stimulation-induced expression of select target genes (*Myc, Irf4, Tnfaip3* and *Ptger4*) was confirmed by RT qPCR (**Figure S2C**). Target genes were largely overlapping, indicating that the two H3ph marks may act coordinately at regulatory regions (H3S28ph) and transcribed regions (H3S31ph) to regulate expression. Importantly, shared target genes (padj < 0.05, Log2 Fold Change > 0.5) are overrepresented in programs involved in GC positive selection^29,30^ and bivalent repressed genes in GC B cells^19^ (**Figure 2A**). Given the important function of PRC2 and H3K27me3 in GC B cells, along with their marked sensitivity to signaling cues and the prominence of transitional chromatin in stimulated cells, we decided to focus on the GC reaction as a relevant model to further investigate this chromatin state. Similar to naïve B cell activation, GC B cell positive selection and fate transitions occur upon interacting with T_FH_ cells through immunological synapses featuring complex sets of ligand/receptor pairs^31,32^ (**Figure 2B**). Of note, exposure of naïve B cells to different immune synapse agonists (anti-BCR, anti-CD40, IL-21 and LPS) for one hour results in H3ph, with individual stimuli resulting in only modest increases and combinatorial stimulation resulting in the highest H3ph levels (**Figure 2C**). This suggests that the quality of signaling information can be directly transduced to chromatin through the deposition of H3ph. In this regard, high immune synapse signaling quality and strength elicit GC B cell positive selection and fate transitions through the rapid shift of the chromatin landscape embedding responsive genes, from a repressed state controlled by PRC2^19,33^ to an activated state dictated by coactivators such as Arid1a^34^, Crebbp^35,36^ and Kmt2d^37,38^. We sought to elucidate how immune synapse signaling cascades trigger this epigenetic switch. To assess immune synapse-dependent histone phosphorylation in GC B cells we used two mouse models (**Figure 2B**): first, the Rosa26-Fucci2aR cell cycle reporter^39^, expressing the fluorescent proteins mCherry and EGFP fused to the cell cycle reporters hCdt1 (G1) and hGem (S/G2/M), respectively (**Figure 2B**), allowed us to gate out mitotic cells featuring high levels of H3ph that could mask immune synapse signaling induced phosphorylation^40,41^. Second, a *cMyc^GFP/GFP^* reporter that harbors the endogenous cMyc protein fused to GFP^42^, and aided the identification of centrocytes undergoing positive selection^43–46^. Rosa26-Fucci2aR mice were immunized with the T-cell dependent antigen sheep red blood cells (SRBCs). Flow cytometry showed the expected frequencies of cell cycle phases, with centroblasts mostly in S/G2/M, and centrocytes mostly in G1 (**Figure 2D**). Immunoblot analyses of sorted centroblasts and centrocytes in G0/G1 at day 9 post-immunization (peak of the GC reaction), showed increased H3S28ph and H3.3S31ph in centrocytes, suggesting induction by selection signals within the light zone (**Figure 2D**). To directly relate stimulation and H3ph, we performed immunoblots on sorted GC B cells from wild type mice stimulated *ex vivo* with anti-IgM, anti-IgG, anti-CD40 and IL-21 for one hour. We found increased phosphorylation levels upon stimulation (**Figure S2D**), indicating that mimicking selection signals through BCR, CD40 crosslinking and IL-21 can rapidly induce H3ph. Next, we immunized *cMyc^GFP/GFP^* mice with SRBC and performed phospho-flow cytometry at day 9. We observed induction of H3S28ph and H3.3S31ph in G1/S Myc-GFP^+^ over G1/S Myc-GFP^-^ GC centrocytes (**Figures 2E and S2E**). We next ascertained the genomic location of these PTMs in GC B cells during an infection featuring robust GC reactions using CUT&Tag for H3S28ph and H3.3S31ph. Wild type mice were immunized with *Plasmodium chabaudi*, and centrocytes and centroblasts were sorted at day 21 after immunization, the peak of the plasmodial induced GC reaction^47^. As newly selected cells are an exceedingly rare fraction within the centrocyte population we compared *ex vivo* stimulated centrocytes with resting centroblasts (which are not receiving any endogenous selection signals). We observed stimulation-dependent H3ph deposition patterns mirroring those detected in stimulated naïve B cells, with H3S28ph covering broad regions and H3.3S31ph delimiting transcribed regions (**Figure 2F**). Differential peak analysis revealed H3ph at genes known to be upregulated upon GC positive selection (**Figures 2G, H; Table S2**), including the transcription factors *Myc*, *Irf4* and *Batf;* the co-stimulatory receptors *Cd86* and *Cd83;* and members of the NF-kB pathway *Nfkbia* and *Nfkbiz*. Importantly, GSEA analysis showed that H3ph target genes are enriched for signatures observed upon immune synapse induced CD40 and NF-kB pathway activation^48^, light zone or centrocyte signatures, and gene sets defining positively selected Myc+ GC B cells^29^ (**Figures 2 I, J; Table S3**). Of note, H3S28ph target genes are enriched for *de novo* bivalent genes in GC B cells, again suggesting the interplay between H3ph and PRC2/H3K27me3. Together, our results indicate that engagement of signaling pathways that promote GC B cell selection in the light zone downstream of the immune synapse receptors CD40, BCR and IL-21R robustly induce H3ph at immune synapse activated genes.

**Figure 2.**
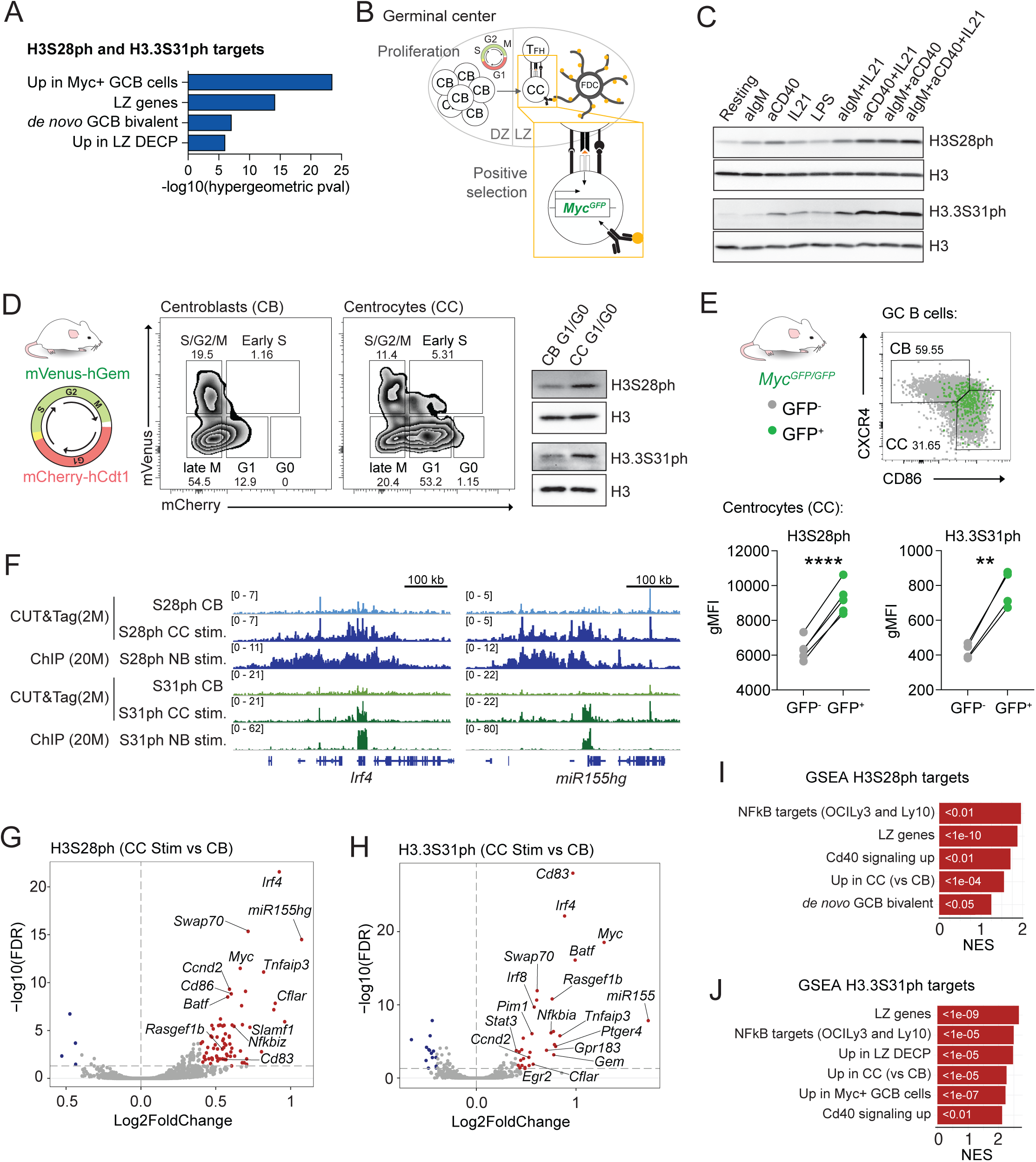
Chromatin integrates signaling quality through the deposition of histone phosphorylation at induced genes upon germinal center B cell positive selection. **(A)** Gene signatures enriched for targets gaining both H3S28ph and H3.3S31ph (padj <0.05, Log2FC >0.5) as determined by hypergeometric mean analysis. **(B)** Schematic of a germinal center with dark (DZ) and light (LZ) zones, proliferative centroblasts (CB) and centrocytes (CC) undergoing positive selection through interactions with follicular dendritic cells (FDC) and T follicular helper cells (TFH) and expression of *Myc* downstream of immune synapse signaling. Two mouse models utilized to assess cell cycle status (FUCCI2aR) and positive selection (*Myc^GFP/GFP^*reporter) are depicted. **(C)** H3S28ph and H3.3S31ph western blots of resting and *ex vivo* stimulated naïve B cells with the indicated stimuli (one-hour). WB representative of two independent experiments **(D)** Schematic of the FUCCI2aR system (left), example of flow cytometry gating (middle) and western blot on the indicated sorted populations (right). Representative of two independent experiments. **(E)** Representative flow cytometry plots highlighting Myc-GFP^+^ cells in germinal center cells (top) and geometric mean fluorescence intensity (gMFI) for H3S28ph and H3.3S31ph in CC (bottom). *n*=4, mean ± s.d. Paired *t* test (two-tailed), ** p-val <0.01, **** p-val <0.0001. **(F)** Example IGV profiles for H3S28ph and H3.3S31ph CUT&Tag and ChIP experiments on CB and *ex vivo* stimulated CC (one-hour anti-IgM, anti-IgG, anti-CD40 and IL-21) and *ex vivo* stimulated naïve B cells (NB) (one-hour with anti-IgM, anti-CD40 and IL21). **(G, H)** Volcano plots of H3S28ph (D) and H3.3S31ph (E) differential deposition in stimulated CC vs CB obtained from CUT&Tag, (*n*=2). **(I, J)** Gene set enrichment analysis (GSEA) for H3S28ph (G) and H3.3S31ph (H) targets, normalized enrichment score (NES), p-values for each signature indicated within the corresponding bar.

### Histone phosphorylation marks chromatin at the time of germinal center B cell selection and directly shapes cell fate transitions

The strong association of H3ph with immune synapse signaling and GC selection-induced programs suggests the PTMs may directly activate chromatin at genes driving GC B cell fate transitions for their rapid and robust transcription. H3ph may activate chromatin through various mechanisms, including altering biophysical properties to increase histone tail mobility and availability to enzymes^49,50^, ejection of polycomb complexes^14,15^, or recruiting chromatin remodeling machinery^8^. To gain insight into the chromatin accessibility changes occurring precisely at the time of centrocyte selection, we performed single-nuclei RNA and ATAC sequencing (10x Multiome) in GC B cells. To increase the proportion of what would otherwise be exceedingly rare selected centrocytes, we immunized *Myc^GFP/GFP^* mice with SRBC, and after 9 days Myc-GFP+ GC B cells were sorted and mixed with total GC B cells (**Figure 3A**). Uniform manifold approximation and projection (UMAP) showed 12 clusters that were annotated by transferring labels from published datasets^51^ and validated previously^34^ (**Figure 3A**). Next, we explored chromatin accessibility across discrete cell fate transitions that include a well-defined population of selected centrocytes. For this, we performed a pseudotime analysis based on expression profiles with initial time anchor at the population of pre-selection centrocytes that just arrived in the light zone (early CC) and final anchor at centroblasts, a population that includes cells that have been selected in the light zone, recycle back to the dark zone and continue to contribute to the GC reaction (**Figures 3B, C**). To validate the pseudotime analysis we plotted gene expression for signatures defining: selected centrocytes upon DEC205-mediated antigen delivery^52^ and dark zone cells (centroblasts)^52^ (**Figure 3C**). Chromatin accessibility at phosphorylated regions along the pseudotime axis revealed that peak chromatin opening precedes expression of programs associated to centrocyte selection. This open chromatin structure decreases in cells that re-acquire dark zone phenotypes (**Figure 3C**). As shown earlier, H3S28ph and H3.3S31ph target multiple genes in common, suggesting that both might be required for their activation; S31ph exerting its activity at promoters and gene bodies, and S28ph at promoters and enhancers. To ascertain whether S28ph is indeed deposited at regulatory regions of S31ph target genes (Log2 Fold Change > 0, p-value < 0.05) we used SCARlink^53^ to identify enhancers from our snATAC/RNAseq dataset (**Figure S3A, Table S4**). Analysis of H3S28ph read density at putative enhancers that could be robustly defined showed increased mean deposition of the mark in stimulated centrocytes compared to centroblasts (and to randomly selected regions not overlapping putative regulatory regions), pointing to a coordinated activity of H3S28ph and H3.3S31ph (**Figures 3D and S3B**). Next, we focused on S31ph target genes that are upregulated upon GC B cell positive selection and analyzed chromatin accessibility at regulatory regions in cell populations before and after receiving immune synapse signaling inputs for selection (pre-selection early CC, selected CC and pre-plasmablasts, which represent a distinct population of selected cells destined for antibody secreting plasma cell maturation), as well as H3S28ph levels in centroblasts and *ex vivo* stimulated centrocytes (**Figure 3E**). We observed low accessibility of H3ph-associated enhancers in early centrocytes and subsequent chromatin opening both in selected centrocytes and pre-plasmablasts, with an increase of H3S28ph at the same regulatory regions in stimulated centrocytes compared to centroblasts (**Figures 3E, F**). Close inspection of selection-associated genes like *Myc* or *Irf4* (**Figure 3G**) showed overlap of H3S28ph peaks with putative enhancers, both in stimulated centrocytes and stimulated naïve B cells. Chromatin accessibility at regulatory regions started increasing upon centrocyte selection and continued to open in early differentiation to plasmablasts. Overall, our results show that H3ph downstream of immune synapse signaling is associated with gained chromatin accessibility and activation of positive selection programs.

**Figure 3.**
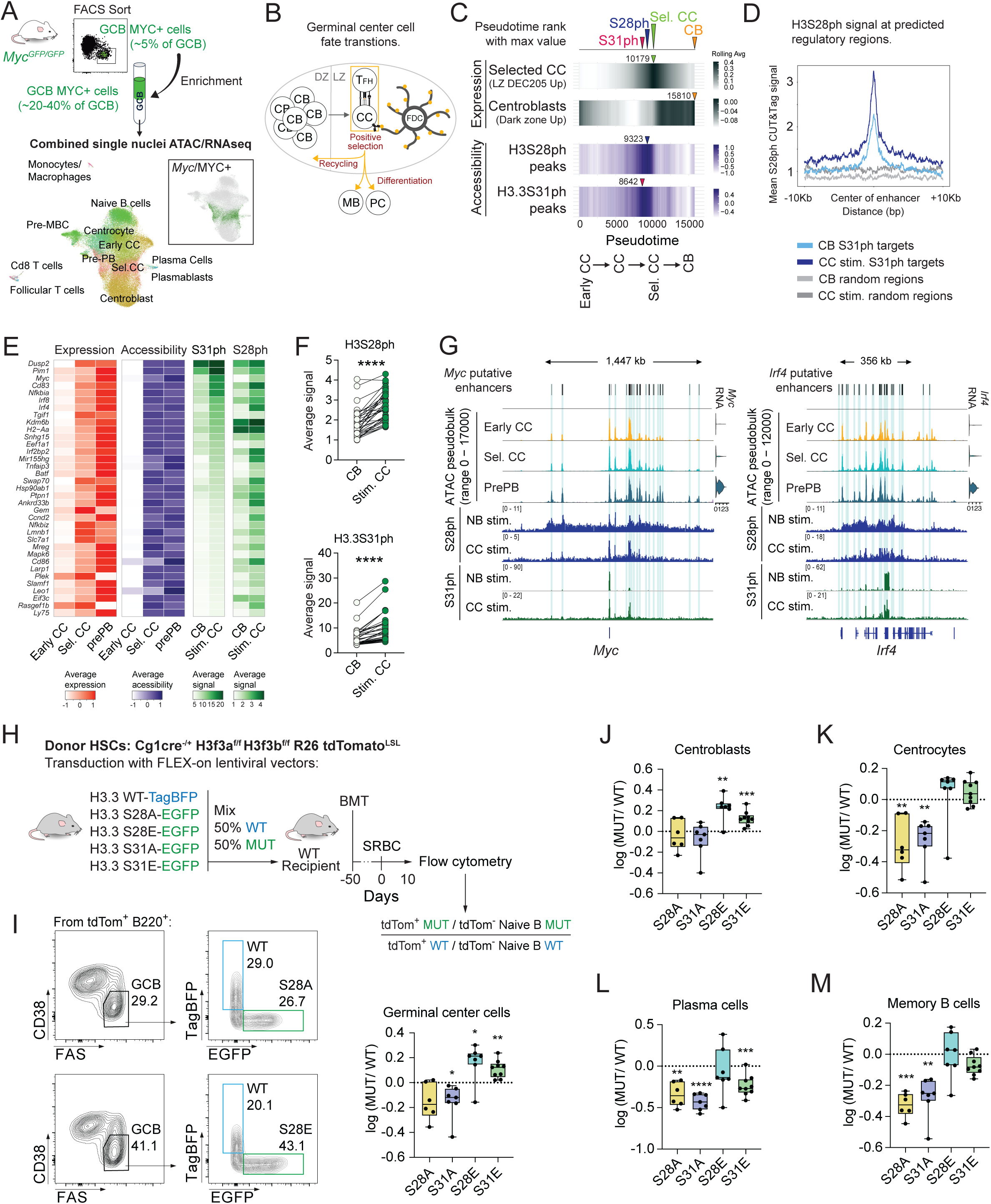
Histone phosphorylation marks chromatin at the time of germinal center B cell positive selection and directly shapes fate transitions. **(A)** Experimental scheme (top) and UMAP of 10x Multiome snATAC/RNAseq on germinal center B cells enriched with selected Myc^GFP+^ cells (bottom). Cell type annotations are based on expression profile datasets. **(B)** Cartoon depicting a germinal center and the different cell fate transitions happening upon positive selection (recycling and differentiation). **(C)** Gene expression profiles (black) defining selected centrocytes and centroblasts, and chromatin accessibility at regions gaining H3S28ph and H3.3S31ph (blue) along a pseudotime performed with initial anchor in early CC and finishing in centroblasts. **(D)** Average H3S28ph CUT&Tag signal over SCARlink-identified regulatory regions of H3.3S31ph target genes and randomly selected regions. (**E**) H3.3S31ph targets, gene expression and chromatin accessibility at SCARlink regulatory regions in early centrocytes, selected centrocytes and pre-plasmablasts; histone phosphorylation status in centroblasts and stimulated centrocytes at gene bodies (H3.3S31ph) and SCARlink regulatory regions (H3S28ph). **(F)** Average CUT&Tag signal for H3S28ph at regulatory regions and H3.3S31ph at gene bodies. Mean ± s.d. Paired *t* test (two-tailed), **** p-val <0.0001. **(G)** Examples showing snATACseq pseudobulk and ChIPseq/CUT&Tag tracks for H3S28ph and H3.3S31ph. (H) *In vivo* GC B cell specific histone genetics experimental scheme. **(I)** Example flow cytometry gates and ratio of frequencies of germinal centers, (**J**) centroblasts, (**K**) centrocytes, (**L**) plasma cell and (**M**) memory B cell populations expressing the indicated histone mutants as a ratio compared with wild type (S28A *n*=6 S31A *n*=7, S28E *n*=7, S31E *n*=9). Mean ± s.d. Paired *t* test (two-tailed), * p-val <0.05, ** p-val <0.01, *** p-val <0.001, **** p-val <0.0001).

Given the potential activity of H3ph histone PTMs in GC B cell positive selection, we aimed to investigate the function of each *in vivo.* Since epigenetic enzymes frequently target multiple residues on histone and non-histone proteins^54^, we sought to directly assess the function of the H3S28 and H3.3S31 residues. For this purpose, we developed a functional histone genetic platform to specifically introduce histone H3.3 point mutants in GC B cells *in vivo* to assess frequencies of GC B cells and differentiated populations. We generated mice with conditional knock out of both H3.3 alleles (*H3f3a^f/f^ H3f3b^f/f^)*, crossed to Cγ1Cre^55^ to delete H3.3 specifically in GC B cells, and to drive a *Rosa26-tdTomato^LSL^* reporter (Cγ1-H3.3-tdTomato). Replacement of the deleted endogenous H3.3 was achieved by transduction of hematopoietic stem cells (HSCs) from Cγ1-H3.3-tdTomato mice with lentiviral particles containing cre-inducible H3.3 wild type or mutant genes that constitutively expressed TagBFP and EGFP, respectively. Serines were mutated to alanines (S28A, S31A) to ablate phosphorylation at these residues; or to glutamic acid (S28E, S31E), mimicking constitutive phosphorylation. Transduced HSCs were expanded^56^ and sorted using TagBFP and EGFP markers. Mixed bone marrow chimeras were generated by injecting a mix of TagBFP-H3.3 WT and each of the mutant EGFP-H3.3 HSCs at a 1:1 ratio into irradiated C57B/6J recipients. After engraftment, mice were immunized with SRBC and splenocytes collected after 10 days (**Figure 3H**). Flow cytometry immunophenotyping was performed, mutant/wildtype ratios of the different population frequencies were calculated and normalized to mutant/wildtype ratio in naïve B cells (before switching endogenous H3.3 expression for transgene expression). Cells expressing the S to A mutant transgenes corresponded with a decreased frequency of GC B cells compared to cells expressing wildtype transgenes (p-value: S28A=0.051, S31A<0.05), while cells mimicking constitutive histone phosphorylation (S to E) had significantly higher frequencies (p-value: S28E<0.05, S31ph<0.005) (**Figure 3I**). Differences in GC B cells were mainly driven by increased frequency of centroblasts in S to E mutants (**Figure 3J**) and decreased centrocyte population in S to A mutants (**Figure 3K**). Plasma cell and memory B cell frequencies of S to A mutant cells were lower than those of wildtype cells (**Figures 3L, M**). On the other hand, S to E mutation did not have an impact in these populations except for the lower frequency observed in S31E plasma cells (**Figures 3L, M**). These *in vivo* GC B cell specific H3.3 mutagenesis experiments reveal the importance of signal transduction to H3.3 serine residues to maintain proper population flux (i.e. recycling vs. differentiation) within the GC reaction. This indicates that phosphorylation of H3.3 serine residues tunes the selection probabilities and fates of GC B cells responding to vaccination.

### Histone phosphorylation antagonizes PRC2 activity via multiple mechanisms

PRC2 mediates *de novo* formation and maintenance of repressed chromatin, often acting across broad domains. PRC2 targets include bivalent genes enriched among immune synapse response genes in GC B cells^19,33^. We hypothesized that expansive PRC2/H3K27me3 is rapidly removed across these loci in response to immune synapse signaling for these genes to be induced during positive selection. We considered that a major mechanism for gene activation by H3ph could be direct antagonism of PRC2 read/write processivity, whereby the enzyme complex both reads (binds) and writes (catalyzes) H3K27me3^57^ and also biophysically condenses chromatin to maintain a feedforward repression loop at its target genes^58^. The robustness of this repression mechanism suggests that active and targeted biophysical mechanisms are likely required to overcome it. It has been previously reported that mitogenic, stress and differentiation signals, as well as direct targeting of the H3S28 kinase MSK1 to the α-globin promoter, correlate with PRC2 displacement, decreased H3K27me3, and gene activation^14,15^. In addition, PRC2 is unable to bind H3K27me3S28ph peptides^14^. Despite these findings, a complete understanding of regulation of PRC2 by H3ph is lacking, in part because PRC2-chromatin contacts extend beyond the H3 tail, locus-wide chromatin repression is complex, and additional mechanisms linking H3ph to de-repression of PRC2 targets are unknown. To fully characterize the interplay between H3ph and PRC2, we designed a series of biochemical and biophysical experiments aimed at addressing multiple PRC2 mechanisms using semi-synthetic nucleosomes and polynucleosomes harboring the native modifications (designer nucleosomes or “dNucs”)^59^. To measure equilibrium dissociation constants for PRC2-nucleosome binding, we performed fluorescence polarization experiments using purified PRC2 core complex (EZH2, SUZ12, EED, and RBBP4) and fluorescein-labeled dNucs. Consistent with a previous report showing a minor role of H3K27me3 on PRC2 binding^60^, we found only slightly higher affinity for H3K27me3 over unmodified mono-nucleosomes which was not statistically significant (*K_d_^app^* 160.8 ± 49.9 nM *vs*. 182.2 ± 26.4 nM; **Figure 4A**). In contrast to the minor effects of H3K27me3 on nucleosome binding, either H3ph significantly disrupted PRC2 binding (*K_d_^app^* of 254.6 ± 53.8 nM for H3S28ph; 275.1 ± 97.1nM for H3.3S31ph: **Figures 4B-C**).

**Figure 4.**
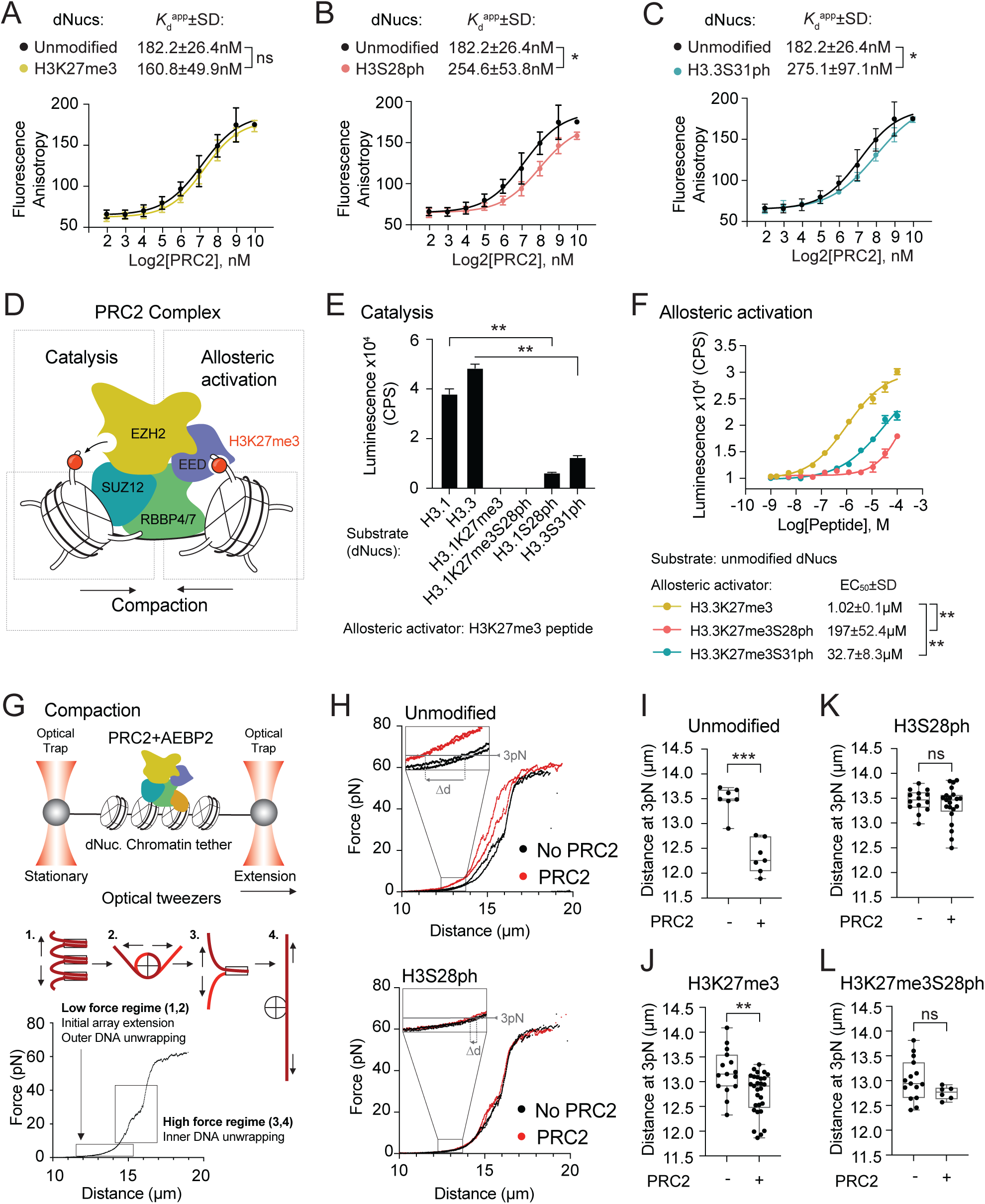
Histone phosphorylation antagonizes PRC2 activity via multiple mechanisms. **(A-C)** Fluorescence anisotropy experiments using a PRC2 tetramer (Eed, Ezh2, Suz12 and Rbbp4) and the indicated designer nucleosomes (dNucs). (Unmodified nucleosomes *n=*5, H3K27me3 *n*=3, H3S28ph *n*=3, H3S31ph *n*=3, mean ± s.d. Unpaired *t* test (two-tailed), * p-val <0.05. **(D)** Scheme of PRC2 activation and catalysis (“read-and-write” mechanism). **(E)** Histone methyltransferase assay to assess PRC2 activity (allosterically activated with H3K27me3 peptides) on the indicated dNuc substrates. **(F)** Histone methyltransferase assay to test PRC2 activity on unmodified (H3.1) nucleosomes upon allosteric activation with the indicated histone tail peptides. (*n*=3, mean ± s.d. Unpaired *t* test (two-tailed), * p-val <0.05, ** p-val <0.01). **(G)** Schematic of the single-molecule optical tweezers experimental setup (top) and a sample force-distance curve (bottom) depicting chromatin parameters analyzed in distinct force regimes (low force range < 5pN; high force range, between 8-40pN). **(H)** Example force-distance curves obtained from unmodified (top) and H3S28ph (bottom) chromatin tethers in the absence or presence of PRC2+AEBP2. Inset describes the force level where the length of different tethers in (I) where measured. **(I-L)** Comparison of chromatin tether extensions (loaded with the indicated designer octamers) at 3 pN of force in the absence or presence of PRC2+AEBP2 complex. Each dot represents a single chromatin molecule (Unmodified *n*=7, unmodified + PRC2 *n*=7, H3K27me3 *n*=15, H3K27me3 + PRC2 *n*=29, H3S28ph *n*=14, H3S28ph + PRC2 *n*=22, H3K27me3S28ph *n*=15, H3K27me3S28ph + PRC2 *n*=6. Unpaired Mann-Whitney *U* test (two-tailed), * p-val <0.05, ** p-val <0.01, *** p-val <0.001).

PRC2 can interact with histone H3 tails through a catalytic pocket located in EZH2 and a regulatory pocket in the EED subunit^61–64^ (**Figure 4D**). PRC2 has minimal basal methyltransferase activity and requires EED-H3K27me3 binding and resulting allosteric changes in the catalytic lobe to enhance activity^61^. Due to the proximity of both S28 and S31 to K27, we wondered if phosphorylation of either residue impacted PRC2 enzymatic activity. To test PRC2 methyltransferase activity we used *trans-*acting H3K27me3 peptides to allosterically activate the complex and detected robust methyltransferase activity towards unmodified (H3.1 or H3.3), and no activity on fully methylated (H3.1K27me3 or H3.1K27me3S28ph) nucleosome substrates (**Figure 4E**). Strikingly, PRC2 activity on phosphorylated nucleosomes (H3.1S28ph or H3.2S31ph) was significantly reduced to near-background levels (**Figure 4E**). In a second series of experiments, we tested the effects of H3ph on PRC2 allosteric activation, a critical feature of the enzyme complex’s read/write mechanism (**Figure 4D**). Here H3.3K27me3, H3.3K27me3S31ph or H3.3K27me3S28ph peptides were used as allosteric activators for PRC2-mediated methylation of an unmodified nucleosome substrate. Of note, either phosphorylation significantly reduced the efficiency of allosteric activation by more than 30-fold, increasing the peptide concentration needed to support PRC2 methyltransferase activity (EC_50_ 1.02 ± 0.1 μM for H3.3K27me3; 197± 52.4 μM for H3.3K27me3S28ph; 32.7 ± 8.3 μM for H3.3K27me3S31ph, **Figure 4F**).

Finally, since PRC2 can compact and stabilize chromatin via biophysical interactions^58,65^, we tested the effect of H3ph on mechanical properties and compaction of PRC2-bound chromatin by performing single-molecule force manipulation assays using a dual-trap optical tweezers instrument (**Figure 4G**). In this assay, a single molecule of biotinylated lambda DNA was tethered between optically trapped beads, and unmodified or modified histone octamers loaded *in situ* using the NapI histone chaperone^66^. Force was then measured while mechanically stretching chromatin tethers in the presence or absence of PRC2 (here a pentamer containing the AEBP2 subunit, which confers stability to the core complex^62^ and stronger nucleosome binding^67^) (**Figure 4G**). We focused analysis at the low force range (< 5 pN), during which neighboring nucleosomes unstack and the outer nucleosomal DNA turn unwraps^68–70^ (**Figures 4G, H**). At 3pN the length of chromatin tethers (loaded with comparable amounts of octamers, **Figure S4A**) containing unmodified and H3K27me3 nucleosomes became shorter upon PRC2 addition, reflecting mechanical compaction by the pentameric complex (**Figures 4I, J**). In contrast, incorporation of H3S28ph to either unmodified or H3K27me3 chromatin abolished PRC2-mediated compaction (**Figure 4K, L**). Mean chromatin tether length difference (correlating to the extent of compaction) with addition of the PRC2 complex was smallest for H3K27me3S28ph and H3S28ph chromatin substrates (**Figure S4B**). H3.3S31ph chromatin, in contrast, showed significant PRC2-mediated compaction, suggesting specific activity of H3S28ph interfering with PRC2-mediated chromatin compaction rather than generic effects of H3ph, such as increase in negative charge (that would also be a feature of H3.3S31ph nucleosomes) (**Figure S4C**). Our results are consistent with a model in which H3S28ph impairs PRC2-dependent chromatin compaction.

This series of biophysical and enzymatic assays establish that both phosphorylation marks weaken PRC2 binding to chromatin, potently disrupt allosteric activation, and render nucleosomal substrates suboptimal for PRC2 methyltransferase activity. Further, H3S28ph is selectively capable of interfering with PRC2-mediated mechanical compaction of chromatin.

### Phosphorylation dependent rapid epigenetic derepression and augmented chromatin interactivity within regulatory domains containing inducible genes

Given these biochemical and biophysical insights into H3ph-mediated antagonism of PRC2 function, we investigated whether H3ph could be mediating the activation of chromatin at PRC2-repressed targets in response to immune synapse signaling. We immunized wild type mice with *P. Chabaudi*, sorted GC B cells and stimulated *ex vivo* with anti-IgM, anti-IgG, anti-CD40 and IL-21 to mimic immune synapse signaling for 0, 1, 3, 6, 18, 36 and 60 hours and performed CUT&Tag. We did not observe substantial differences between 0 hours and the early time points (1, 3 or 6 hours; not shown). However, starting at 18 hours we detected a clear decrease of H3K27me3 at H3ph target genes, including *Batf* and *Ccnd2* (**Figure S5A, Table S2**). Given the domain-level deposition of both H3K27me3 and H3S28ph, we focused on H3K27me3 regions gaining H3S28ph in stimulated centrocytes (Log2 Fold Change > 0, p-value < 0.05) and performed differential analysis (**Figure 5A**). Genes showing decreased H3K27me3 upon stimulation (18 hours) included known PRC2 repressed immune synapse induced genes, including known bivalent genes^19^ (e.g. *Nfkbie, Irf4, Myc, Egr2, Sub1, Satb1*). Closer inspection of individual genes (**Figures 5B and S5B**) uncovered extensive overlap of H3K27me3 regions with H3S28ph domains, and regions that lose methylation upon stimulation coincide with those gaining H3S28ph, strongly suggesting that stimulation-induced deposition of H3S28ph along large chromatin domains could activate swaths of PRC2-repressed chromatin harboring regulatory regions to promote transcription. In line with this notion, we observed that regulatory regions linked to the indicated genes (by SCARlink) are contained within domains that gain H3S28ph and lose H3K27 methylation upon receiving signaling inputs (**Figure 5B**). To test if those regulatory regions become activated in response to stimulation, we performed CUT&Tag on H3 lysine 27 acetylation (H3K27ac) stimulated GC B cells following conditions used previously (**Table S2**). Differential analysis of H2K27ac peaks overlapping H3S28ph domains (Log2 Fold Change > 0, p-value < 0.05) showed a preferential shift towards increased deposition (**Figure 5C**). Additionally, H3K27ac deposition is negatively correlated to that of H3K27me3 at phosphorylated regions (**Figure S5C**, p-value = 5.19 e-09), indicating that domains gaining H3S28ph undergo an epigenetic switch permissive for transcriptional activation (**Figure 5A-C**, **S5C**). Indeed, H3K27me3 and H3K27ac peaks overlapping phosphorylated domains undergo significantly more loss and gain of these marks, respectively, than those not overlapping (**Figure 5D**).

**Figure 5.**
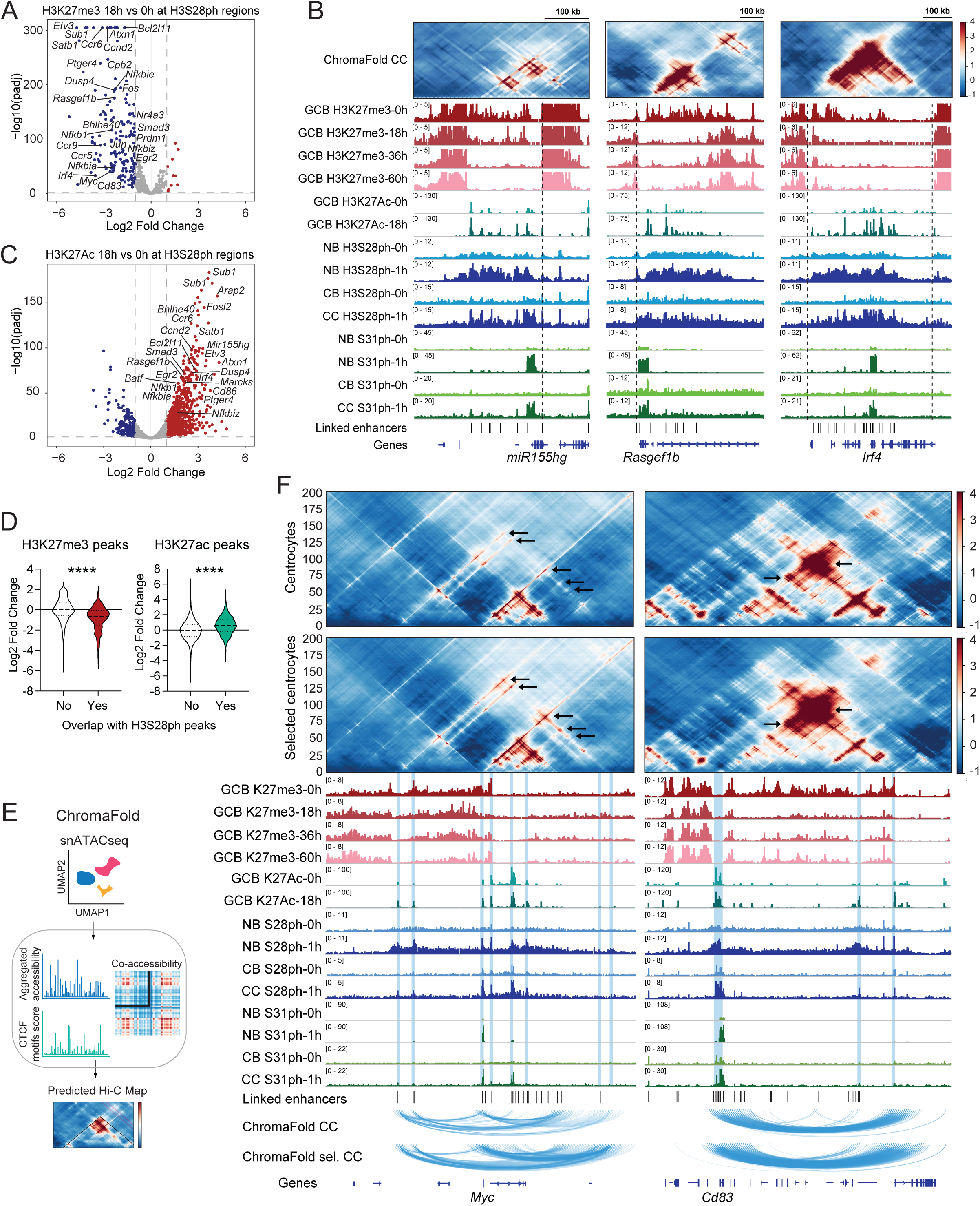
Phosphorylation dependent rapid epigenetic derepression and augmented chromatin interactivity within regulatory domains containing inducible genes. **(A)** H3K27me3 differential deposition (CUT&Tag) in germinal center B cells 18 h after stimulation (anti-IgM, anti-IgG, anti-CD40 and IL-21) compared to 0 h. Regions depicted are peaks overlapping H3S28ph domains gained in stimulated CC (*n*=2). **(B)** Example ChromaFold prediction of chromatin 3D interactions in centrocytes (CC), IGV profiles for the indicated ChIP and CUT&Tag experiments in germinal center B cells (GCB), naïve B cells (NB), centrocytes (CC) and centroblasts (CB) at the indicated stimulation time points (conditions as in A), and SCARlink predicted regulatory regions linked to the specified genes. **(C)** H3K27ac differential deposition (CUT&Tag) in germinal center B cells (conditions as in A). (*n*=2). **(D)** H3K27me3 and H3K27ac log2 Fold Changes of peaks overlapping and not overlapping with H3S28ph peaks. Unpaired Mann-Whitney *U* test (two-tailed), **** p-val <0.0001). **(E)** ChromaFold schematic to predict chromatin contact maps from single cell accessibility data. **(F)** Examples of ChromaFold predictions in centrocytes and selected centrocytes and IGV profiles for H3K27me3, H3K27ac, H3S28ph and H3.3S31ph (CUT&Tag/ChIP) in germinal center B cells (GCB), naïve B cells (NB), centrocytes (CC) and centroblasts (CB) before and after stimulation. SCARlink predicted enhancers linked to *Myc* and *Cd83*. Arcs at the bottom show discrete interactions as predicted by ChromaFold.

Genes and their regulatory regions are usually located within topologically associating domains (TADs). We wondered whether H3ph would counteract PRC2 activity within TAD boundaries, thus favoring intra-domain chromatin interactivity to promote enhancer contacts and induce target gene expression. To test this, we used ChromaFold^71^, a deep learning model to predict chromatin 3D contacts from single cell accessibility data (**Figure 5E**). We used our combined single cell ATAC/RNA-seq dataset with enrichment for Myc^+^ GC B cells (selected centrocytes) (**Figure S5D**) to obtain Z-score normalized chromatin contact maps from centrocytes and selected centrocytes. We detected 5,011 TADs and close inspection of individual genes revealed that H3S28ph deposition is often delimited by TAD boundaries (**Figure 5B top panels**). We observed statistically significant differences in 2,360 TADs, with a majority displaying gained interactivity upon selection (**Figures S5E, F; Table S5**), including TADs containing light zone induced genes such as *Ccnd2*, *Myc* or *Cd83*. Notably, TADs gaining interactivity were also associated with 78 and 38 genes targeted by H3S28ph and H3.3S31ph in stimulated centrocytes, respectively (**Figures S5E, F; Table S5**). A closer inspection of two highly histone phosphorylated genes, *Myc* and *Cd83*, (**Figure 5F**) showed that H3S28ph deposition was delimited by TAD boundaries in centrocytes, displaying a clear overlap with H3K27me3 domains that lose methylation upon stimulation. Increased interaction frequency in selected centrocytes occurred preferentially at regions gaining H3S28ph, H3.3S31ph and H3K27ac (**Figure 5F** black arrows, blue arcs), with a strong association between H3S28ph deposition and gained chromatin interactivity. Thus, H3ph at PRC2-repressed domains is associated with H3K27me3 loss, H3K27ac gain with increased chromatin interactivity, and induced transcription.

### Phosphorylation dependent Polycomb domain derepression is a general feature of rapid gene induction across immune cells

To test if the deposition of H3S28ph is a broadly employed mechanism for derepression of Polycomb domains, we stimulated naïve CD4 T and NK cells as described previously and performed CUT&Tag 24-hours post-stimulation. Differential analysis of H3K27me3 deposition in CD4 T cells shows preferential methylation loss at H3S28ph regions containing genes required for T cell differentiation (*Spry1*, *Tbx21*, *Nfatc1*, *Egr2*) (**Figure 6A, Table S6**). As in GC B cells, H3K27me3 loss preferentially occured within H3S28ph domains (**Figure 6B**). Similarly, in NK cells we observed loss of H3K27me3 at phosphorylated domains encompassing genes upregulated upon activation and required for NK maturation and survival (*Gpr171*, *miR155hg*, *Il2ra*, *Il15ra*) (**Figure 6C, D, Table S6**). Our results suggest that domain-level depostion of H2S28ph is a commonly utilized mechanism to enable rapid signaling-dependent derepression of PRC2 target loci.

**Figure 6.**
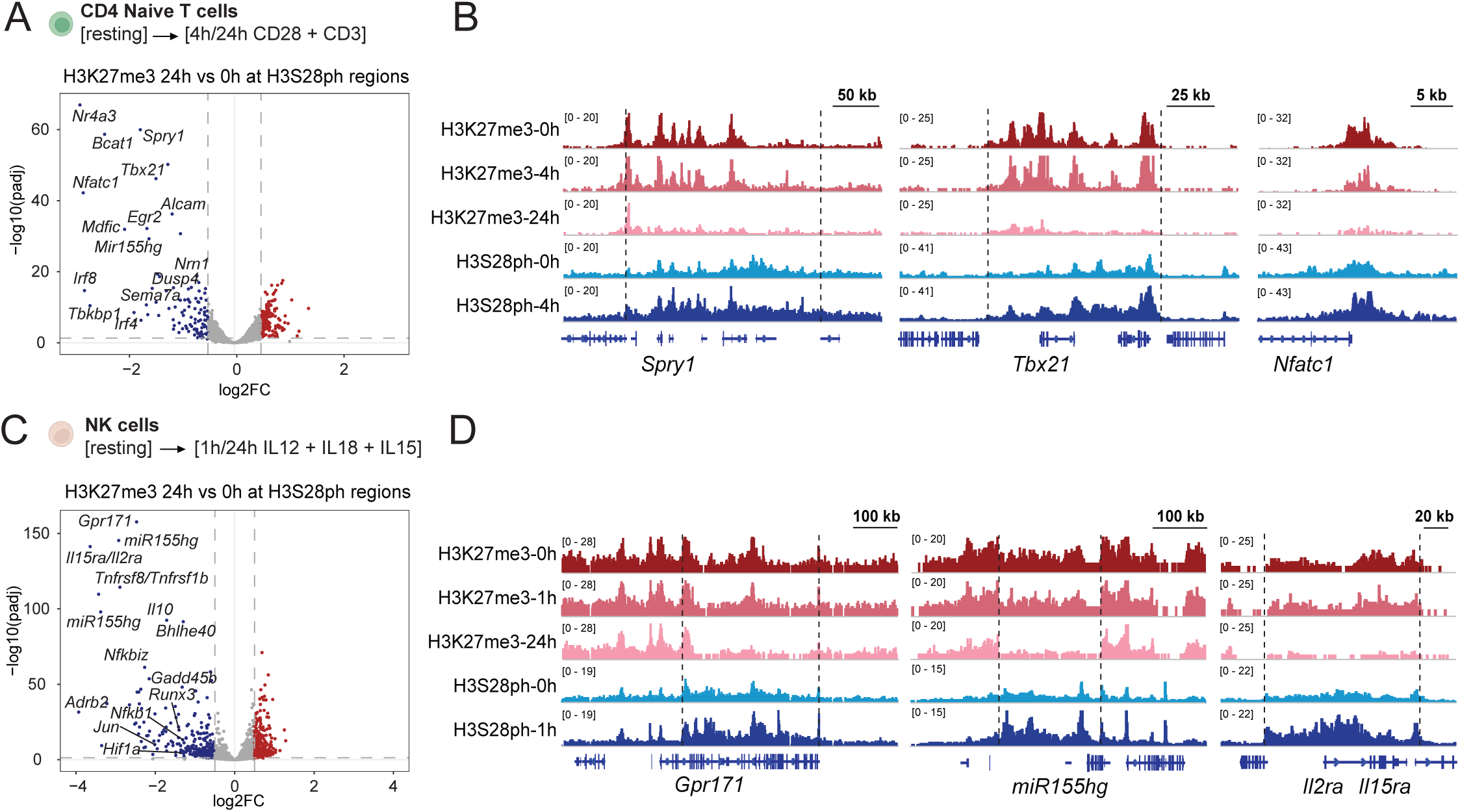
Phosphorylation dependent Polycomb domain derepression is a general feature of rapid gene induction across immune cells. **(A)** Differential analysis of H3K27me3 CUT&Tag peaks in resting and stimulated CD4 T cells, showing only peaks overlaping regions gaining H3S28ph after stimulation and **(B)** example IGV tracks for H3K27me3 and H3S28ph in resting and stimulated naïve CD4 T cells. **(C)** Differential analysis of H3K27me3 CUT&Tag in resting and stimulated natural killer (NK) cells and **(D)** example tracks.

### Phosphorylation-dependent derrepressed programs are maintained in an activated state through deposition of lysine 36 methylation at histone H3

Histone phosphorylation is a transient PTM, with a fast deposition (15-30 minutes) and lasting up to 3-4 hours^9,10^. We hypothesized that H3S28ph acts as initial barrier for PRC2 activity, thus allowing early transcription; its activity could be coordinated with another more stable mechanism to prevent PRC2 from re-methylating the template. Di-methylation of H3 at lysine 36 (H3K36me2) is a broad histone modification with well-established roles opposing Polycomb activity both biochemically and *in vivo*^72–74^. To ascertain whether H3K36me2 increases at regions harboring H3S28ph, we performed H3K36me2 CUT&Tag in steady state and stimulated GC B cells as previously described. Individual examples showed higher H3K36me2 levels 18-hours after activation, coincident with H3K27me3 loss and H3K27ac gain, and with a deposition strikingly similar to that of H3S28ph (**Figures 7A**, **S6A**). This strongly suggests the interplay between H3K36me2 and H3S28ph potentiates and maintains activated chromatin domains, which is also supported by the negative correlation (p-value = 2.75 e-11) between H3K36me2 and H3K27me3 peaks overlapping H3S28ph regions (Log2 Fold Change > 0, p-value < 0.05) (**Figure 7B**). Genes embedded in the activated domains include genes required for memory cell differentiation (Ccr6^75^) and plasma cell differenciation (Sub1^76^), pointing to the need of H3K36me2 to keep expression of those genes in fate comitted cells.

**Figure 7.**
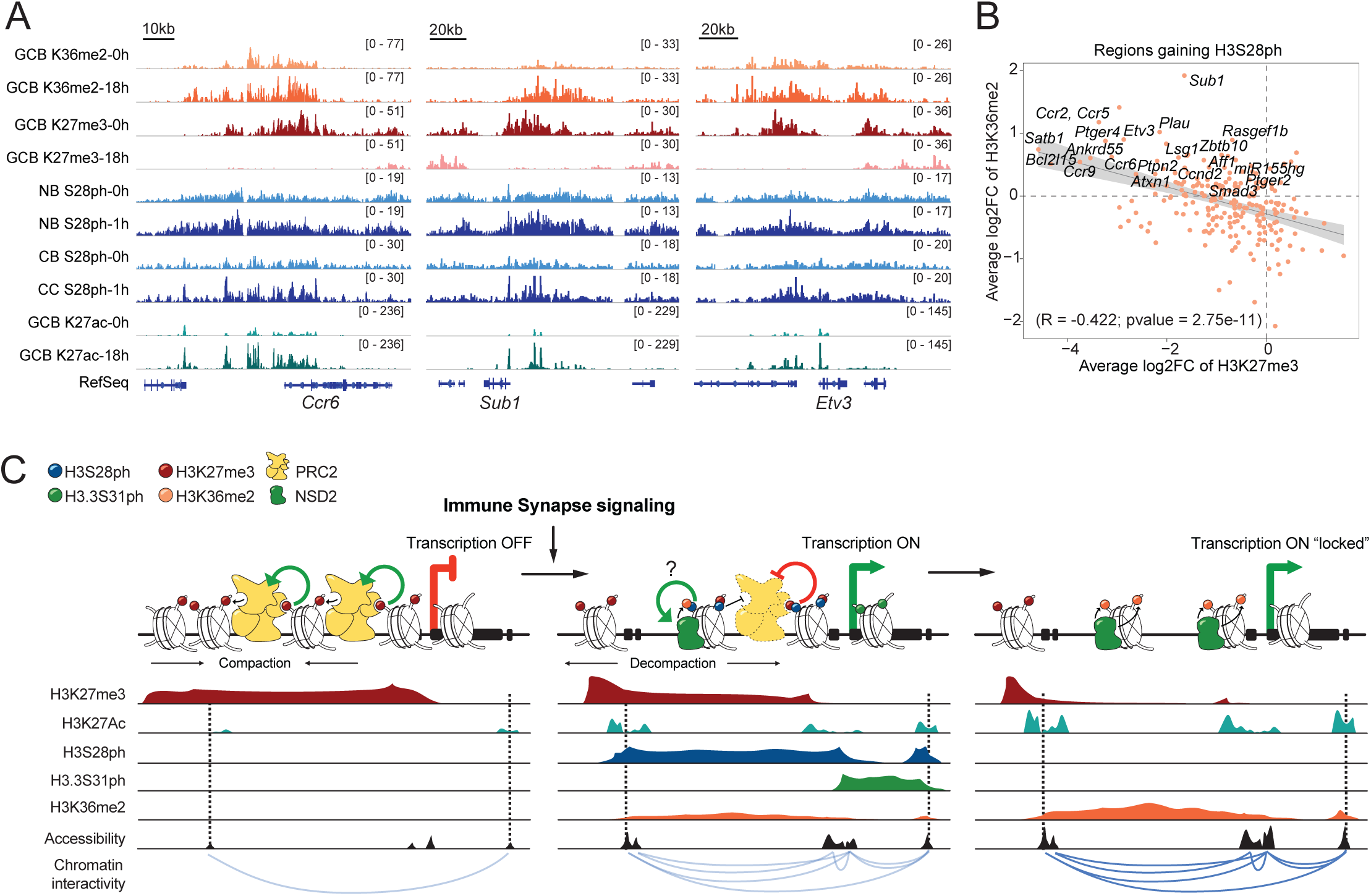
Phosphorylation-dependent derrepressed programs are maintained in an activated state through deposition of lysine 36 methylation at histone H3. **(A)** Example IGV tracks for the indicated histone posttranslational modifications in steady state and stimulated B cell populations. **(B)** Plot depicting average log2 fold change of H3K27me3 and H3K36me2 peaks overlapping peaks of increased H3S28ph after stimulation of CC. **(C)** Model describing signaling-induced chromatin de-repression: signaling responsive genes are embedded within chromatin domains repressed by PRC2/H3K27me3. Immune synapse signaling results in deposition of H3S28ph and H3.3S31ph, that impairs PRC2 read/write mechanisms, decreases H3K27me3, and promotes chromatin decompaction, enhancer activation (H3K27ac) and increased regulatory interactivity to favor transcriptional activation. Transient H3S28ph could favor activity of the K36 methyltransferase NSD2 to keep chromatin domains and contained loci in a stable transcriptionally active state.

Together, our data establish that immune synapse signal-induced H3ph activates chromatin by disrupting PRC2-mediated repression leading to decompaction, enhancer activation, increased intra-domain dynamics, and transcription activation critical for controlling GC B cell fate decisions (**Figure 7C**). We propose a model in which transiently deposited H3S28ph favors H3K36me2 catalysis to maintain chromatin in a stable active state to promote distinct cell fates (**Figure 7C**).

## Discussion

Here, we discover a common stimulation-induced chromatin state characterized by the presence of repressive H3K27me3 and signal-dependent H3S28ph at critical activation and fate determining genes across multiple primary cell types. Further, we define mechanisms of H3ph-mediated activation of these repressed domains for rapid signaling-induced transcription and revel physiologic contexts of phospho-methyl chromatin switches. While feedforward mechanisms governing chromatin repression, such as those employed by PRC2, have been extensively characterized, insight into dedicated mechanisms that enable the selective derepression of developmentally regulated or stimulation-induced genes has been missing. Stereotypic of this, germinal center B cell fate transitions are subject to a delicate balance between chromatin repressors and coactivators that must be rapidly shifted in response to immune synapse signaling inputs to drive selection and fate determination. Tight control is required to allow GC B cells to proliferate while undergoing immunoglobulin somatic hypermutation, affinity-dependent selection, and discrete fate transitions, all processes critical for the generation of B-cell mediated immunity. In general, it is not known how repressive signals can be rapidly reversed, and how such epigenetic “switches” are triggered by signaling inputs. We demonstrate that B cell selection and differentiation signals lead to H3S28ph and H3.3S31ph deposition at regulatory regions and gene bodies. We demonstrate that H3ph represents an initial epigenetic event — the transduction of signaling inputs into the chromatin template at select target genes — and coordinates a cascade of chromatin de-repression and activation to regulate inducible transcription. Moreover, phosphorylation levels increase with cumulative signaling inputs, suggesting that chromatin integrates diverse immune receptor signaling inputs through histone phosphorylation to translate signaling quality into probabilities of chromatin activation at lineage defining loci thereby controlling cell fate transitions. We describe an epigenetic switch involving the reactivation of PRC2 repressed targets, including GC B cell bivalent genes^19,33,77^, and establish a sequence of events from immune synapse signaling to histone H3 phosphorylation to PRC2 antagonism and regulatory chromatin activation, enabling the rapid induction of GC B cell selection programs. Our comprehensive characterization of the interplay between PRC2 and histone phosphorylation builds onto previous findings linking H3S28ph with PRC2 function^14,15,78^ and establishes the potent inhibitory effect of H3.3S31ph and H3S28ph and their association with PRC2 removal from promoters and distal regulatory regions in selected centrocytes. Importantly, we establish the importance of this rapid epigenetic switch for physiologic cell state transitions through defining its mechanistic and functional activity in the response to immunization and infection.

Our studies also reveal previously unknown functions for the two H3ph PTMs, and highlight their distinct activities in different, mostly non-overlapping, chromatin contexts. Within the GC reaction, H3.3S31ph throughout the transcription unit may reinforce promoter derepression of certain targets (i.e. *Ccnd2*, **Figure S5A**), where there is clear overlap of H3.3S31ph and PRC2/H3K27me3. H3.3S31ph could limit PRC2 activity at promoters and transcribed regions acting together with other modifications such as H3K4me3 and H3K36me3^72,79^. Distinct from H3.3S31ph, H3S28ph overlaps H3K27me3 across large chromatin domains spanning promoters and distal regulatory regions, the activation of which is required to achieve transcription of target genes. Importantly, phosphorylation of H3 at either S28 or S31 in K27me3-bearing nucleosomes interferes with the PRC2 EED-EZH2 auto-activating feedback loop that is a defining feature of PRC2’s robust repression activity. Therefore, signaling-dependent deposition of H3ph rapidly antagonizes PRC2 activity, and facilitates the subsequent demethylation of chromatin domains (**Figure 5B**). Future work will establish the involvement of active demethylation by KDM6A/UTX or KDM6B/JMJD3, or passive loss due to nucleosome turnover inherent to regulatory regions^80^ or cell proliferation^45^.

The avidity of PRC2 for chromatin depends on the additive effect of multiple low affinity interactions, such as those with histone tails, and a strong electrostatic engagement with DNA^60,64,81^. The effect of H3S28ph on PRC2 bound chromatin could be due to the combination of (i) diminished affinity of EED and EZH2 for the histone tail, (ii) competition of H3ph with DNA for binding to the positively charged PRC2 surfaces, thereby impairing PRC2 binding to DNA at the nucleosomal exit^81^ and looping and bending of linker DNA^82,83^. Interestingly, the decompacting effect of S28ph on H3K27me3 chromatin suggests that initial S28ph deposition could facilitate access of additional transcription factors or other machineries, even while H3K27me3 is still present. Supporting this hypothesis, phosphorylated regions in selected centrocytes are associated with chromatin opening, enhancer activation (augmented H3K27ac deposition), increased chromatin interactivity and expression of target genes. We suggest that while this rapid activation requires the disruption of PRC2 function, additional H3ph dependent mechanisms could act synergistically, including the induction of p300/CBP catalytic activity and enhanced H3K27ac deposition^9,11,17^ or recruitment of 14-3-3 proteins and chromatin remodeling complexes^8^. In this regard, our results suggest an additional H3S28ph dependent mechanism — the increased deposition of H3K36me2 — that may participate in stabilizing or locking in the active state through long-term antagonism of PRC2^72–74^ following the transient early H3S28ph mediated effects. Further investigation relating to this model is required, including whether increased H3K36me2 is directly impacted by H3S28ph or indirectly increased due to loss of H3K27me3/PRC2. Given the strikingly similar deposition of H3S28ph and H3K36me2, is tempting to speculate that phosphorylated chromatin could promote NSD2 activity (the primary K36 di-methyltransferase in GC B cells^84,85^) by stabilizing binding or favoring catalytic activity (**Figure 7C**). Mechanisms described here also have implications in disease states. For instance, transient polycomb derepression is sufficient to induce malignant transformation in *Drosophila*^86^, and highly aggressive GC derived Activated B Cell Diffuse Large B Cell Lymphomas (ABC-DLBCL) harbor mutations that promote the constitutive activity of immune synapse signaling pathways^87^.

Functional studies of histone PTMs have been challenged by multi-copy histone genes and complexities of the enzyme systems and their targeting of multiple histone and non-histone residues. To directly assess the function of specific H3 residues *in vivo* we established a novel genetic histone replacement platform. With this approach, we demonstrated that both H3S28ph and H3.3S31ph are required to maintain appropriate GC B cell fate transitions and amplify selection and recycling. While exhaustive study of these models is required, we propose that mutants abrogating residue-specific histone phosphorylation (*e.g.*, H3S28A), mimic low signaling inputs, while phosphomimetic serine to glutamic acid (S to E) mutants mimic higher signaling strength/quality, associated with increased positive selection.

We propose a model in which chromatin integrates the quality of diverse cell signaling events through specific histone phosphorylations. This signal scaffold activity of chromatin then initiates a cascade of epigenetic events, including PRC2 inhibition, co-activator stimulation, and broad domain activation and mobility, thus promoting specific regulatory DNA interactivity and the activation of local gene transcription. Our study uncovers a potent mechanism for rapid activation of repressed domains via histone phosphorylation, generalizable to diverse contexts including the initiation of developmental transcription programs and the induction of stimulation responsive genes.

## Data and code availability

Multiome: 10.5281/zenodo.14164741, 10.5281/zenodo.14164927, 10.5281/zenodo.14165393, 10.5281/zenodo.14165809, GSE237990, GSE249081. H3.3S31ph and H3K36me3 ChIP in naïve B cells: 10.5281/zenodo.13901056. H3S28ph_ChIP in naïve B cells: 10.5281/zenodo.13900120. H3S28ph, H3.3S31ph, H3K27me3 and H3K4me3 CUT&Tag in GC B cells: 10.5281/zenodo.13882304. H3K27ac, H3K36me2 CUT&Tag in GC B, NK and CD4 T cells datasets and H3K27me3, H3K27ac, H3K4me1, H3K4me3 and H3.3 CUT&Tag in naïve B cells: 10.5281/zenodo.14719444 and 10.5281/zenodo.16739394. Macrophages CUT&RUN and HiC: 10.5281/zenodo.16794971. HiC in naïve B cells: GSE143852. H3ph ChIPs in Macrophages: GSE63792 and GSE125159. Optical tweezer data collection custom script: 10.5281/zenodo.14166360.

## Acknowledgements

This work was supported in part by R01AI148416, AAI Intersect Award (S.Z.J.), Lymphoma Research Foundation Fellowship (A.M.P.), STARR-I14-0020 (S.Z.J., C.S.L., A.M.M., S.L.). F30CA275379 and T32GM152349 (J.T.C.). National Science Foundation Graduate Research Fellowship grant 2139291 (M.K). P01CA272295-01A1 (S.Z.J., W.B., A.M.M., L.C., C.R.F., M.R.G., C.E.M.).

## Author contributions

Conceptualization: S.Z.J., A.M.M., A.M.P. and W.B. Biochemical, epigenomic, single cell and mouse experiments: A.M.P., W.B, A.W.D., M.J.B., C.J., D.J.A., A.R., V.E.K., I.K. NK cells experiment: M.O. and A.M.P. Binding assays: J.C. supervised by Y.L. Methyltransferase assays: M.R.M., L.K., B.G., C.E.S. supervised by M.-C.K. Optical tweezers: J.C., A.M.P. and R.S. supervised by S.L. Computational analysis: Y.S., M.K., R.Y. and W.W. supervised by C.S.L. Also A.M.P., C.R.C., M.J.B. and A.M.T. Writing – Original Draft: A.M.P and S.Z.J. Writing – Review & Editing: all authors.

## Supplementarty Materials

**Table S1**: H3S28ph and H3.3S31ph ChIPs differential analysis related to Figure S2.

**Table S2**: H3S28ph, H3.3S31ph, H3K27me3 and H3K27ac CUT&Tag differential analysis related to Figures 2, 5 and 7.

**Table S3**: GSEA analysis related to Figure 2.

**Table S4**: SCARlink final putative enhancers related to Figure 3.

**Table S5**: Differential analyses of ChromaFold predicted Topologically Associating Domains related to Figure 5.

**Table S6**: H3K27me3 differential analysis related to Figure 6.

## Methods

### Mouse models

All experimental procedures performed with laboratory mice followed institutional guidelines and were approved by the Research Animal Resource Center and Institutional Animal Care and Use Committee (IACUC) of Weill Cornell Medicine. All mice utilized in this study were between 8-14 weeks of age. Both females and males were used in all experiments, except for C57B/6J recipient mice that were all males.

H3f3a^f/f^ H3f3b^f/f^ mice were kindly gifted by Laura Banaszynski. Rosa26-Fucci2aR was developed by J. Jackson group^39^. Remaining strains were obtained from The Jackson Laboratory: C57B/6J (CD45.2, stock 000664), Cγ1-Cre (B6.129P2(Cg)-Ighg1tm1^(cre)Cgn^/J) (stock 010611), Rosa26Stop-tdTomato (AI14, stock 007914), Myc^GFP^ (stock 021935).

### Immunizations

To induce the formation of splenic germinal centers (GC), mice were immunized by intraperitoneal injection with 500 µl of 2% sheep red blood cells (SRBCs; Cocalico Biologicals 20-1334A) diluted in sterile 1x Phosphate-buffered saline (PBS). For experiments requiring higher numbers of GC B cells, animals were immunized with *Plasmodium chabaudi*. Parasites were maintained as frozen stocks and passaged in C57B/6J wild type mice. Parasitemia was assessed by flow cytometry after staining 1-2 µl of blood with Hoechst33342 (Invitrogen) and anti-Ter119 (eBiosciences). Mice were immunized by intraperitoneal injection with 10^5^ *P. Chabaudi*-infected erythrocytes from the donor mouse.

### Hematopoietic stem cell expansion, transduction and bone marrow chimera generation

Hematopoietic stem cells (HSC) were obtained by collecting bone marrow cells from tibias and femurs from donor strains, performed red blood cell lysis and stain with antibodies Ly-6A/E (Sca-1)-Pacific Blue, Biolegend 108120, Cd117 (c-kit)-PE/Cy7, Biolegend 105814, biotinylated lineage markers TER-119 79748, B220 79752, Cd11b 79749, Cd3e 79751, Ly6G/Ly6C 79750 from Biolegend). Lineage negative, Sca-1 positive and c-Kit positive cells were sorted, resuspended in polyvinyl alcohol (PVA) HSC expansion media as previously described^56^ and plated in human fibronectin coated 96-well plates (Corning BioCoat 354409). Media was replaced every other day and transductions performed at expansion day 6. For viral particle production for transductions, HEK293T cells (ATCC CRL-3216) were transfected using Lipofectamine 3000 (Thermo Scientific L3000015) following manufacturer recommendations. Vectors used were psPAX2, pMDG.2 and different pLV(FLEXon)-EGFP vectors containing the H3.3 point mutant versions H3.3S28A, H3.3S28E, H3.3S31A, H3.3S31E or pLV(FLEXon)-TagBFP encoding wild type H3.3 (Vector Builder). EGFP and TagBFP are constitutively expressed from these vectors, while H3.3 transgenes are cre dependent. Supernatants were collected the next 2 days and used for transductions. At transduction day one, 300ul of viral suspension was added to each well of a 96/well fibronectin coated plate and centrifuged for 1 hour at 2000g. Media was discarded, and HSCs were added to the wells, and spun for 10 minutes at 450g. Media was removed and 300 ul of viral suspension with 8ug/mL polybrene (Millipore Sigma TR-1003-G) were added, plate was spun for 1 hour at 1000g and subsequently transferred to 37°C to incubate for 2 hours. Media was replaced for PVA expansion media. Next day media was discarded and viral supernatant with polybrene was added. Plate was centrifuged for 1 hour at 1000g and incubated and 37°C for 2 hours. Media was replaced and cells were expanded for 6 days. Transduced cells were sorted based on EGFP of TagBFP and were kept expanding for a few more days. For bone marrow chimera generation, transduced HSCs were injected through the retro-orbital sinus into lethally irradiated C57B/6J males (two 450 rad doses on a RS 2000 Biological Research X-ray Irradiatior, Rad Source Technologies) and immunophenotyping performed after 6-8 weeks.

### Splenocytes flow cytometry and cell sorting

Splenocyte cell suspension was obtained by disaggregating tissue, filtering through 100 um and 40 um cell strainers and performing Ficoll (Cytiva 17-5446-02) gradient separation of splenic live mononuclear cells. In case of immunizations with *P. Chabaudi*, splenocyte cell suspension was treated with ACK lysis buffer for removal of red blood cells. Cells were subsequently blocked with Fc block (Biolegend 101302), stained (antibodies below) and analyzed by flow cytometry on BD Fortessa or BD Symphony S6 analyzers. When needed, cells were fixed and permeabilized using the BD Cytofix/Cytoperm Kit (554714, BD Biosciences). For cell sorting B-cells were enriched by negative selection (Miltenyi Biotec 130-090-862) or germinal center cells with anti-peanut agglutinin antibodies (Miltenyi Biotec 130-110-479). Cells were sorted on BD FACS Aria II or BD Symphony S6 sorters. Antibodies and viability markers used are as follows: antibodies from BD Biosciences: B220-PE/Cy7 552772, B220-FITC 553087, B220-BV786 563894, CXCR4-Biotin 551968, CXCR4-PE 561734, IgD-BV510 563110, FAS-BUV805 741968, FAS-PE/Cy7 557653, FAS-BV421 562633, CD138-BUV737 564430, CD38-BUV395 740245. Antibodies from BioLegend: CD86-PE/Cy7 105014, CD86-BV605 105037, IgD-APC 405714, TCRb-APC/Cy7 109220, CD3-APC/Cy7 100222, CD19-APC/Cy7 115530, NK1.1-PE 108708, CD49b-PacBlue 108918, Streptavidin-APC/Cy7 405208, Streptavidin-APC 405207, zombie NIR viability 423106 and DAPI 42280. From eBioscience CD38-APC 17-0381-81 and Fixable Viability Dye-eFluor 506 from Thermo Fisher Scientific 65-0866-14.

### *Ex vivo* stimulations

Bone marrow derived macrophages were obtained by collecting bone marrow from tibias and femurs, peform RBC lysis and plate in 1:3 L929 conditioned medium and 2:3 complete RPMI (Corning 10-040-CM) ff10% FBS (Peak), Glutamax (Gibco 35050061), penicillin/streptomycin (Gibco 15140122), Hepes (Gibco 15630080), non-essential amino acids (Gibco 11140050) and 2-mercaptoethanol (Gibco 21985023)]. After 10 days, media was replaced with complete RPMI with 2.5 ng/ml IL-3 and 2.5 ng/ml M-CSF (PreproTech 213-13 and 315-02) for 4 days. Cells were then stimulated for one hour with lipopolysaccharides from Salmonella typhosa (Sigma-Aldrich 2387). Splenocyte suspensions were prepared as described in the previous section. Naïve B cells were enriched by negative selection (Miltenyi Biotec 130-090-862), resuspended in complete RPMI. Cells were incubated for 20 min at 37°C in a humidified atmosphere of 5% CO2 and then treated with 10 µg/ml anti-CD40 (Thermo Fisher Scientific 16-0402-86), 10 µg/ml anti-IgM (Jackson ImmunoResearch 115-005-020) and 5 ng/ml IL-21 (R&D Systems 594-ML-010) for the indicated time. GC B cells were sorted, resuspended in complete RPMI and treated with the same mix as above but with addition of 10 µg/ml anti-IgG (Thermo Fisher Scientific 16-5098-85) during the indicated time points. For long culture time points (36 and 60 hours), viable cells (DAPI negative) were sorted before performing subsequent experiments. Naïve CD4 T cells were enriched by negative selection (Miltenyi Biotec 130-104-453), resuspended in complete RPMI with 10ng/ml IL-2 and 10ng/ml IL-7 (PreProtech, 212-12 and 217-17). Cells were incubated for 3 hours, IL-7 removed and then stimulated for 4 and 24 hours with 10ng/ml IL-2 and plate-bound 1ug/ml CD28 and 5ug/ml CD3 (BioXCell, BE0015-1 and BE0001-1). NK cells were enriched (Miltenyi Biotec 130-115-818) and sorted, resuspended in complete RPMI and stimulated for 1 and 24 hours with 20 ng/ml IL-12 (R&D systems 419-ML-050), 10 ng/ml IL-18 (Mblbio B002-5) and 5 ng/ml IL-15 (Miltenyi Biotec 130-095-760).

### Hi-C

Macrophages were prepared as described in previous section and HiC performed using the HiC+ kit (Arima genomics). HiC-Pro (3.1.0) was used to align the individual Hi-C replicates to mm10, with alternative contigs removed. Only valid ligation products and reads passing a quality threshold of MAPQ30 were retained, but otherwise default settings for all HiC-Pro settings were maintained. The final merged libraries were obtained by combining all *.allValidPairs files where duplicates across biological replicates are maintained. These files were then processed with Juicer (1.19.01) to produce Hi-C files containing resolutions 5 and 25 Kb.

### Immunoblot

Cells were washed once with iced-cold PBS and pelleted to perform lysis in 2x laemmli buffer (50ul/1M cells) with boiling for 5 minutes. Proteins were resolved in 4-20% Tris-glycine gels (Thermo Fisher Scientific XP04205BOX) and transferred to PVDF membranes (Bio-Rad 1620177). Incubation with primary antibodies was performed overnight at 4C, and secondary antibodies were incubated for 1 hour at room temperature. Antibodies are anti-H3S28ph (Abcam 32388), anti-H3.3S31ph^10^, anti H3 (Cell signaling technologies 9715), HRP-conjugated anti Rabbit IgG (GE NA9340V or DAKO P0399). Imaging was performed in an iBright system (Thermo Fisher Scientific) after incubation with chemiluminescent substrates (Protein biology 34580 and 34095).

### RT qPCR

RNA was isolated using RNAeasy Kit (Qiagen). For RT–PCR, extracted RNA was treated with DNase and cDNA was synthesized using High-Capacity cDNA Reverse Transcription Kit (Applied Biosystems). Quantitative PCR (qPCR) was performed using SYBR green dye (Applied Biosystems) and normalized to *Gapdh*. Primer sequences as follows: Myc-F: GCAGCGACTCTGAAGAAGAGC, Myc-R: GGATGGAGATGAGCCCGACT, Tnfaip3-F: GGAAAGCCAGAAGAAGCTCA, Tnfaip3-R: ATGAGGCAGTTTCCATCACC, Ptger4-F: ACCATTCCTAGATCGAACCGT, Ptger4-R: CACCACCCCGAAGATGAACAT, Irf4-F: GGAGCCAGCTGGATATCTCT, Irf4-R: CTGGCAGAGAGCCATAAGGT, Gapdh-F: TGGTGAAGGTCGGTGTGAAC, Gapdh-R: TCATCCACCTCCCCACAGTA.

### Chromatin immunoprecipitation

ChIPs were performed as previously^9^ using antibodies described in immunoblot section.

### CUT&Tag/RUN

Genomic mapping by CUT&Tag was performed using CUTANA pAG-Tn5 (EpiCypher) as per recommendations (EpiCypher protocol v1.7). Antibodies for histone phosphorylation marks are the same used for immunoblotting. Other antibodies used are H3K27me3 (Cell signaling technologies 3737), H3K27ac (Thermo Fisher MA5-23516), H3K36me2 (Epicypher RD130056), IgG (Abcam 46540) and secondary anti-rabbit IgG (Antibodies Online ABIN101961), secondary anti-mouse IgG (Abcam 46540). Primers for library amplification and indexing are described previously^88^.

### ChIP and CUT&RUN/Tag analysis

Samples were processed using an in-house nextflow-based pipeline (https://github.com/michaelbale/jlabflow). Briefly, reads were trimmed using the Cutadapt wrapper trim-galore and mapped to mm10 with bowtie2 using *--very-sensitive-local* with a maximum insert size of 1000 bp. The initial bam file was then filtered for a minimum MAPQ of 30. Subsequently, PCR duplicates and reads mapping to the mitochondrial chromosome or within regions form a unified forbidden list from ENCODE and^89,90^ were eliminated. Finally, only correctly mated mapped reads were retained. Peak calls were made using EPIC2^91^ on each individual sample with *--bin-size/--gaps-allowed* adapted to the different histone modifications (H3K27me3 and H3S28ph 500/5; H3K36me3 and H3.3S31ph 300/5, H3K4me3 200/3). Peak overlaps were determined using the bedtools^92^ intersect function with default parameters. For NK and CD4 T cells, overlapping regions marked by H3S28ph and H3K27me3 were analyzed by calculating fold changes in the normalized H3S28ph signal (stimulated vs. unstimulated). Regions within the top 20th percentile of fold change were classified as showing increased signal. For broad histone modification data (H3S28ph and H3K27me3), sequencing depth normalization factors were calculated using DESeq2^93^ estimateSizeFactors on count matrices of 1000 bp flanking regions on either side of each peak (any overlap of flanking and peak set was removed using bedtools subtract). The flanking normalized sizeFactors were used to compute bigwigs and perform differential analysis. Differential peak analysis was carried out using DESeq2 and all peaks with an FDR ≤ 0.05 were called as differential. Peaks were annotated to overlapping genes by using ChIPseeker version 1.34.1^94^ annotatePeak. Genes were ranked by the Log2FC value and run GSEA using MSigDB database and curated gene signatures from literature, and only gene sets with adjusted p-value of 0.05 were considered. Hypergeometric distribution was used to determine the extent to which a pathway was overrepresented in the list of common H3S28ph and H3.3S31ph target genes for **Figure 2**.

### ChromHMM state discovery and annotation

ChromHMM^5^ software was used to binarize normalized epigenetic signal assaying H3S28ph, H3.3S31ph, H3K27ac, H3K27me3, H3K4me3, H3K4me1, and H3.3 in bone marrow derived macrophages (resting and one-hour stimulated) and naïve B cells (resting and one-hour stimulated) in 200 bp bins. Data was binarized and chromHMM 12 state discovery was performed independently for each cell type, stacking resting and stimulated data for discovery across treatment within each cell type. States in both models were annotated based on emission parameters, overlap enrichment with genomic annotation (TSS, gene body, etc) and gene sets for responsive genes in stimulated BMDMs and naïve B cells, and heuristic assessment of genome segmentation.

### 10x Multiome Single cell RNA and ATAC sequencing

Nuclei were isolated from sorted germinal center B cells according to the ‘Low Cell Input Nuclei Isolation’ protocol (10x Genomics CG000365-Rev B) and processed using Chromium Controller & Next GEM Accessory Kit (10x Genomics 1000202) and Chromium Next GEM Single Cell Multiome <u>ATAC</u> + Gene Expression Reagent Kit (10x Genomics 1000285) following manufacturer’s User Guide (10x Genomics CG000338-Rev D). Single-cell RNA and <u>ATAC</u> sequencing libraries were prepared using Dual Index Kit TT Set A (10x Genomics 1000215) and Single Index Kit N Set A (10x Genomics 1000212) respectively and sequenced on an Illumina NovaSeq6000 platform.

### Multiome analysis

Multiome data was processed as previously described^34^. Briefly, multiome fastq files were processed using CellRanger Arc version 2.00 (10x Genomics). CellRanger outputs were processed using Signac version 1.5.0^95^. Peaks were called using MACS2^96^ with default settings. Samples were combined to generate a combined peak set from all samples. Peaks larger than 10kb and less than 20 bases were filtered from this peak set. Harmony version 0.1.0^97^ was used on PCA based on RNA and LSI based on ATAC to correct batch effect between experiments. Signac FindMultiModalNeighbors was used on these Harmony corrected values to generate multimodal nearest neighbors upon which the UMAP is based. Cell type labels were assigned based on RNA expression using the Seurat TransferLabel function and reference datasets^51^. Gene signature module scores for expression were calculated using the AddModuleScore function, with a control value of 5. Accessibility module scores were calculated using the AddChromatinModule function using BSgenome.Mmusculus.UCSC.mm10 as the reference genome. Early centrocytes (EarlyCC) were defined as cells expressing intermediate levels of light zone and dark zone markers, *Myc* negative and negative for Myc+ and LZ_DECP signatures. Selected CC (SelCC) were defined by combining two clusters form the label transfer: the CC_Rec cluster which contains cells positive form Myc transcript, and Recycling cluster which contains cells positive for Myc+ and LZ_DECP signatures. From these cells, Pre-plasmablasts were defined as cycling *Bcl6*-low non-pre-memory cells. Slingshot version 2.8.0^98^ was used to generate pseudotime based on the first and second Harmony corrected PCA of expression, with the starting anchor being EarlyCC and final anchor being dark zone cells (centroblasts). Rolling averages of 500 cells across pseudotime were calculated for both gene expression and chromatin accessibility scores. Heatmaps were generated by extracting average expression and accessibility from the DotPlot function for the indicated genes and linked peaks. Peaks and genes were linked using SCARlink as detailed in the next section. Code for multiome processing can be found at https://github.com/crchin/multiome_process.

### SCARlink putative enhancer identification

SCARlink^53^ gene-level models were trained using all cells that had paired scATAC and scRNA data from the filtered 10x multiome object for H3.3S31ph target genes (p-value < 0.05, Log2FC > 0 in H3.3S31ph CUT&Tag centrocytes vs centroblasts). Models were trained on scATAC accessibility tiles from 750 kb upstream to 750 kb downstream of gene bodies for non-sparse genes (scRNA sparsity < 95%). Shapley analysis on cell type annotations identified tiles considered to be putative enhancers in CC_Rec (z-score > 0.5, FDR < 0.001) for each gene-level model with a Spearman correlation of greater than 0.05 (between predicted expression and true expression). *Cd86* (Spearman correlation = 0.0408) links were included only for Figures 3E and 3F based on heuristic measurement of acceptable model performance. Background tile sets were randomly sampled from all tiles that were not putative enhancers in recycling centrocytes to approximate the distribution of distance to gene (on a per-gene level) as the putative enhancers had. Both tile sets (putative enhancers and background tiles) were merged to combine adjacent tiles mapped to the same gene, and duplicate tiles mapped to different genes were merged to be one region mapping to multiple genes. Deeptools^99^ computeMatrix was used to generate matrices of normalized H3S28ph CUT&Tag from centroblasts and centrocytes (2 replicates of each) in a 100kb window with 100bp bins centered at each tile in both tile sets to plot the average profile across replicates.

For heatmaps on histone phosphorylation (**Figure 3**) Deeptools multiBigwigSummary was used to calculate normalized signal in gene bodies and SCARlink putative enhancers for H3.3S31ph and H3S28ph, respectively, in replicates of centroblasts/centrocytes. H3.3S31ph signal was averaged across replicates and plotted. H3S28ph signal was averaged across all enhancers per gene, then averaged across replicates and plotted.

### Hi-C contact map prediction and differential analysis

All cells were clustered using scRNA-seq into UMAPs, with only B cells selected for Hi-C contact map predictions. Specifically, we compared the scATAC-seq and Hi-C contact map between the centrocytes and selected centrocytes. Selected centrocytes are a combination of three clusters from the label transfer: CC_Rec, Recycling (both defined in Multiome analysis) and CB_Rec_Sphase (the latter includes selected cells that are transitioning to the dark zone: still express the Myc+ signature, have already entered S phase and start to express dark zone program). scATAC-seq data were preprocessed following instructions from the ChromaFold GitHub repository. Chromatin co-accessibility was calculated using the ArchR R package^100^, and Hi-C contact maps were predicted using ChromaFold^101^, which was trained using HiC-DC+ normalized contact maps as the target. HiC-DC+^102^ is an R package that normalizes Hi-C contact maps by fitting a negative binomial regression model that accounts for genomic distance, GC content, mappability and effective bin size. This allows us for direct use of the z-scores to identify significant interactions in downstream analysis. Interactions up to 2Mb from the diagonal on the contact map were predicted. Predicted Hi-C contact maps from ChromaFold were further quantile normalized to eliminate batch effects. To evaluate TAD differences around specific genes under different conditions, we assessed the intensity of interactions around the gene and used the Wilcoxon Rank-Sum test and Kolmogorov-Smirnov (KS) test to evaluate differences between the conditions. P-values were adjusted for multiple testing, and the average intensity change between conditions was calculated. We further overlapped the differential TADs with H3.3S31ph and H3S28ph targeted genes. TADs were identified as significantly different if their adjusted p-value from the Wilcoxon Rank-Sum test was less than 0.1 and Log2FC was greater than 0.5. We identified 78 differential TADs overlapped with S28ph targeted genes, and 38 TADs for S31ph targeted genes (padj <0.1 and Log2 Fold Change > 0.5).

### Protein Expression and Purification

Recombinant four- and five-subunit PRC2 complexes were expressed and purified as previously^103^. The four-subunit PRC2 core complex contains EZH2, SUZ12, EED and RBBP4 proteins, all with a PreScission protease cleavable N-terminal 6xHis6-MBP (hexahistidine-maltose binding protein) tags. The five-subunit PRC2 complex additionally contains AEBP2 with a PreScission cleavable Strep-II tag. Expression vectors based on the pFastBac plasmid system (Invitrogen) were transformed into dH10Bac (Invitrogen) competent cells. Following manufacturer guidelines, baculovirus stocks (P0, P1 and P2) were generated in sf9 insect cells grown in Sf-900 III SFM medium (Invitrogen). P2 titer was determined by a commercially available service based on gp64 expression (Expression Systems). 1L of Tni insect cells (Expression Systems) were seeded at 2×10^6^ cells/mL in ESF-921 Protein Free Medium (Expression Systems) and infected at a M.O.I of 1.0 with each baculovirus. Cultures were grown at 27 °C, 130 rpm for 66 hours before cells were harvested, snap-frozen, and stored at −80 °C until purification.

For complex purification, frozen Tni cells were lysed in Lysis Buffer (10 mM Tris pH 7.5, 250 mM NaCl, 0.5% IGEPAL CA-630, 1mM TCEP) treated with 1x Pierce^TM^ protease Inhibitor (Thermo Fisher). Cell debris was removed by centrifugation at 29000 rcf for 40 minutes. Clarified lysate was incubated with amylose resin (NEB) equilibrated in Lysis Buffer for 2 hours under gentle rotation. Lysate-bead mixture was added to a glass 2.5cm x 20cm Econo-Pac Chromatography Column. The column was washed with 10 column value (cv) of Lysis Buffer, followed by 16 cv of high salt buffer (10 mM Tris pH 7.5, 500 mM NaCl, 1 mM TCEP), and 16 cv of low salt buffer (10 mM Tris pH 7.5, 150 mM NaCl, 1 mM TCEP). Protein was eluted off the column in 3 cv of Elution Buffer (Wash Buffer 2, 10 mM Maltose). Elution was collected across five fractions. Relevant fractions were combined and concentrated in Amicon Ultra-15 Centrifugal Filter Unit, 30kDa MWCO. The NaCl concentration was increased to ~250 mM and PreScission was added with a mass ratio of 1 part in 50 for overnight cleavage of MBP tags. The protein was then loaded onto a HiTrap Heparin HP 5 x 5 mL (Cytiva) affinity column equilibrated in Start Buffer (10 mM Tris pH 7.5, 150 mM NaCl, 1 mM TCEP). The protein was eluted in a linearly increasing gradient of Elution Buffer (10 mM Tris pH 7.5, 2 M NaCl, 1 mM TCEP) across 35 cv at flow rate of 1.5 mL/min. Relevant fractions were combined and concentrated as above, then subsequently loaded onto a HiLoad 16/600 Superdex 200pg SEC column (Cytiva) equilibrated in Sizing Buffer (20 mM HEPES pH 7.5, 150 mM NaCl, 1 mM TCEP). Protein was eluted in Sizing Buffer at a flow rate of 0.5 mL/min. Relevant fractions were combined and concentrated as above. The protein was divided into single-use aliquots, which were then snap-frozen and stored at −80 °C until use.

### Nucleosome reconstitution

Nucleosomes were reconstituted using histone octamers (EpiCypher) and 200-bp 5’-FAM-labeled DNA containing the Widom sequence (underlined below)^104^: 5-FAM-TAAAACGACGGCCAGTGAATTGCCGCATCG<u>GAGAATCCCGGTGCCGAGGCCGCTCAATTGGTCGTAGA</u> <u>CAGCTCTAGCACCGCTTAAACGCACGTACGCGCTGTCCCCCGCGTTTTAACCGCCAAGGGGATTACTC</u> <u>CCTAGTCTCCAGGCACGTGTCAGATATATACATC</u>GATTGCATGTAGCTTGGCGTAATCATGGTC Nucleosome reconstitution was performed as previously^105^. In brief, DNA was mixed with histone octamers in three separate ratios (1:0.8, 1:1 and 1:1.2) in histone refolding buffer (2 M NaCl, 6 mM Tris pH 7.5, 0.3 mM EDTA and 0.3 mM TCEP). The mixture was first incubated at 37 °C for 30 min, and then the following volumes of reconstitution buffer (20 mM Tris pH 7.5, 1 mM EDTA, 1 mM dithiothreitol) were added in 30-min intervals: 10.8, 12, 28 and 64 µl. Nucleosome assembly quality was assessed by electrophoresis in a native 6% polyacrylamide 1XTBE gel stained with Redsafe, and the best DNA/histone octamer ratio used for downstream binding assays.

### Fluorescence Polarization

Binding affinities between PRC2 and mononucleosomes were determined by fluorescence polarization. Various concentrations (4 nm - 1 μM) of PRC2 complex were prepared by serial dilutions in binding buffer (50 mM Tris pH 7.5, 10 mM KCl, 0.1 mM ZnCl2, 2mM BME, 0.1 mg/mL BSA, 5% glycerol). An equal volume of FAM-nucleosome (5 nM final in binding buffer) was added, and binding reactions incubated at room temperature for 30 minutes before Fluorescence Anisotropy measurements (TECAN Spark instrument). Here FAM fluorophore was excited at 485 nm with a bandwidth of 20 nm (+/- 20 nm), and emission measured at 535 nm with a bandwidth of 25 nm. Anisotropy values were plotted against protein concentration, with a standard binding curve generated for each nucleosome with baseline adjusted to the anisotropy value of the control sample without protein. Apparent dissociation constant *K*_d_^app^ and Hill coefficient were calculated in *GraphPad Prism* 10 from the mean value of three independent replicates.

### Methyltransferase assays

PRC2 methyltransferase activity with diverse substrates and allosteric activating peptides was measured using the bioluminescent methyltransferase Glo (MTase-Glo; *Promega*). For substrate comparisons, in a 384-well microplate we combined 2.5 μL nucleosome (400nM; *EpiCypher*), 2.5 μL mixture of H3_ff13-24]_K27me3 allosteric activator (120 μM), PRC2 complex (100 nM; *Active Motif*), 3 μL S-Adenosyl Methionine (SAM) and 2 μL MTase-Glo reagent (stock diluted 8x in H_2_O before use). The plate was sealed, briefly centrifuged, and shaken at 450 rpm for 1-2 minutes prior to incubation at 23°C for 2 hours. Reactions were quenched and incubated for 30 minutes with 10 μL of MTase-Glo detection reagent (diluted 4x in H_2_O before use).

For allosteric activator peptide comparisons, we combined 2.5 μL allosteric activator (titration series as noted), a 2.5 μL mixture of H3.1 unmodified nucleosome (400nM) and PRC2 complex (100 nM), 3 μL SAM and 2 μL MTase-Glo reagent (stock diluted 8x in H_2_O before use). The plate was sealed, briefly centrifuged, and shaken at 450 rpm for 1-2 minutes prior to incubation at 23°C for 2 hours. Reactions were quenched and incubated for 30 minutes with 10 μL of MTase-Glo detection reagent (diluted 4x in H_2_O before use).

Luminesence was measured on a 2104 *PerkinElmer EnVision* reader using standard optics and settings. Data was gathered in duplicate or triplicate and analyzed using *GraphPad Prism* 10’s four parameter logistic non-linear regression model.

### Single-molecule experiments

Single-molecule experiments were performed at room temperature on a dual-trap optical tweezers instrument (LUMICKS C-Trap). A computer-controlled stage enabled rapid movement of the optical traps within a five-channel microfluidic flow cell. Force data were collected at 50 kHz and 15 Hz. A custom script was utilized to automate the data acquisition of the single-molecule experiments. First, under constant laminar flow, 4.35-μm streptavidin-coated polystyrene beads (Spherotech) were caught in each trap in channel 1 with a trap stiffness of ~0.33 pN/nm. The traps were then moved to channel 2, which contained biotinylated λ DNA (48.5 kbp). By moving one trap back and forth along the axis of the flow direction, a DNA tether could be formed and detected via a change in the force-extension curve. The traps were then moved to channel 3, and the presence of a single DNA tether was verified by the force-distance curve with the flow turned off. Following verification of a single tether, the λ DNA was extended to 12.5 μm (~ 0.5 pN) and then moved to channel 4, which contained 3 nM histone chaperone NapI and ~2 nM of the histone octamer of interest in HR buffer (30 mM Tris acetate pH 7.5, 20 mM magnesium acetate, 50 mM KCl, and 0.1 mg/mL BSA). The DNA tether was incubated with octamers in channel 4 to form nucleosomes until a specific force threshold was reached or until 30 seconds had elapsed. Incorporation of octamers into the DNA tether causes a length reduction due to DNA wrapping and increases force readings. Force increment at loading is directly proportional to the number of octamers loaded. The measured force on the tether before the tether was moved from channel 4 was recorded and used as a proxy for octamer loading. Only tethers with comparable force readings at loading were used. The DNA tether was then moved back to channel 3, which contained HR buffer and 0.25 mg/mL salmon sperm DNA (to assist removing improperly wrapped histone octamers on the DNA tether). The DNA tether was relaxed to 10 μm in channel 3 and then moved to the experimental channel 5. Channel 5 contained reaction buffer (50 mM Tris-HCl pH 7.5, 10 mM KCl, 0.1 mM ZnCl_2_, 1 mM DTT, 2% glycerol, and 0.1 mg/mL BSA) with or without 100 nM PRC2 complex. After a 10-second incubation in channel 5, the forces were set to zero and a force-distance curve was collected, which extended the tether at a speed of 0.25 μm/s until the force reached ~60 pN. Data were collected with this experimental workflow (in the presence or absence of PRC2) for the different octamers containing unmodified H3, H3K27me3, H3S28ph, H3K27me3S28ph and H3.3S31ph.

---

**Figure S1.**
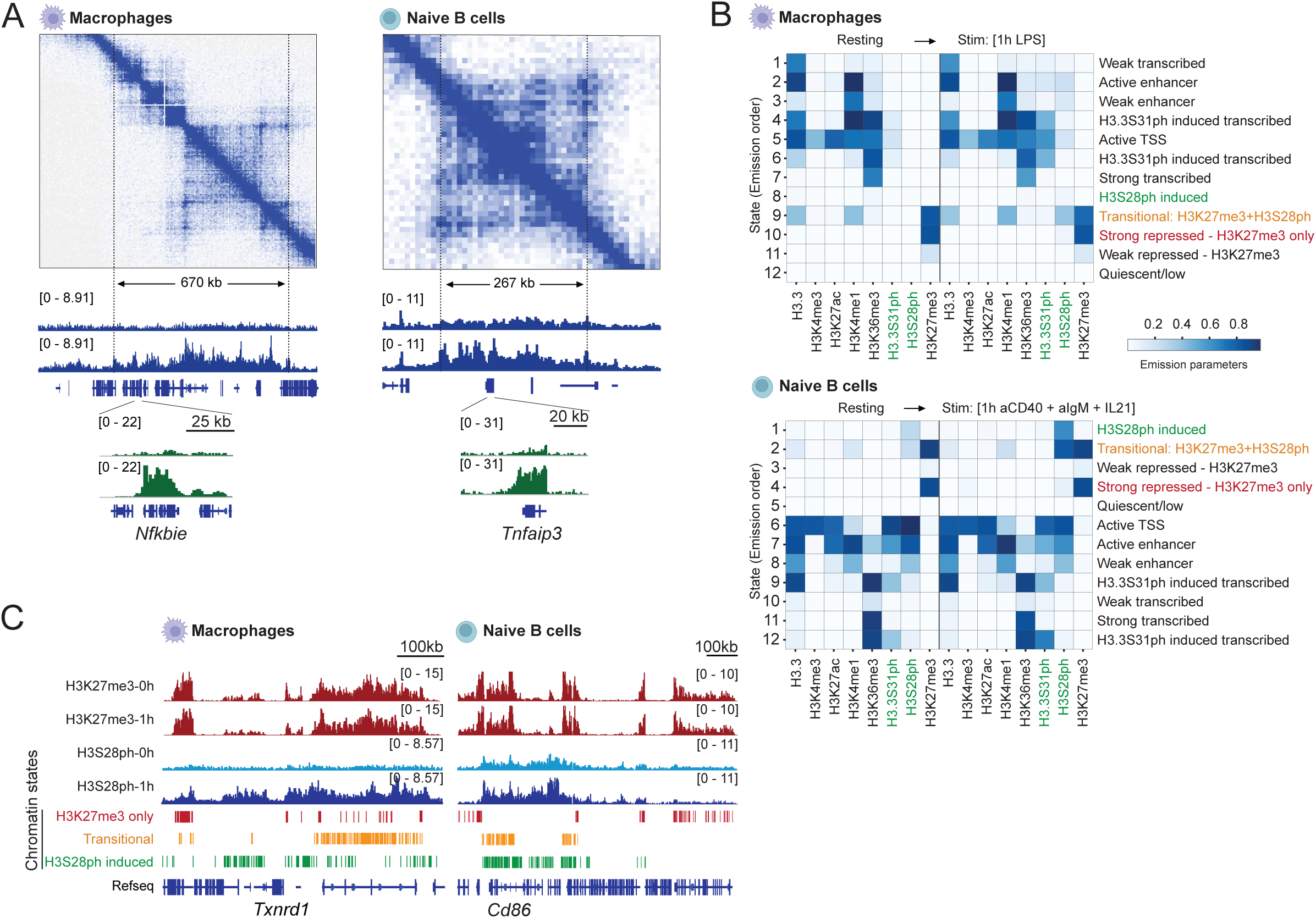
**(A)** Top: HiC contact matrix around *Nfkbie* in resting bone marrow derived macrophages and *Tnfaip3* in naïve B cells. Bottom: example Integrated Genomic Viewer (IGV) tracks from H3S28ph and H3.3S31ph ChIP experiments in resting and stimulated macrophages and naïve B cells. **(B)** Chromatin states discovered by ChromHMM in resting and stimulated macrophages (top) and naïve B cells (bottom). **(E)** IGV tracks for H3K27me3 and H3S28ph and highlighted chromatin states detected around *Txnrd1* in resting and stimulated macrophages and *Cd86* in resting and stimulated naïve B cells.

**Figure S2.**
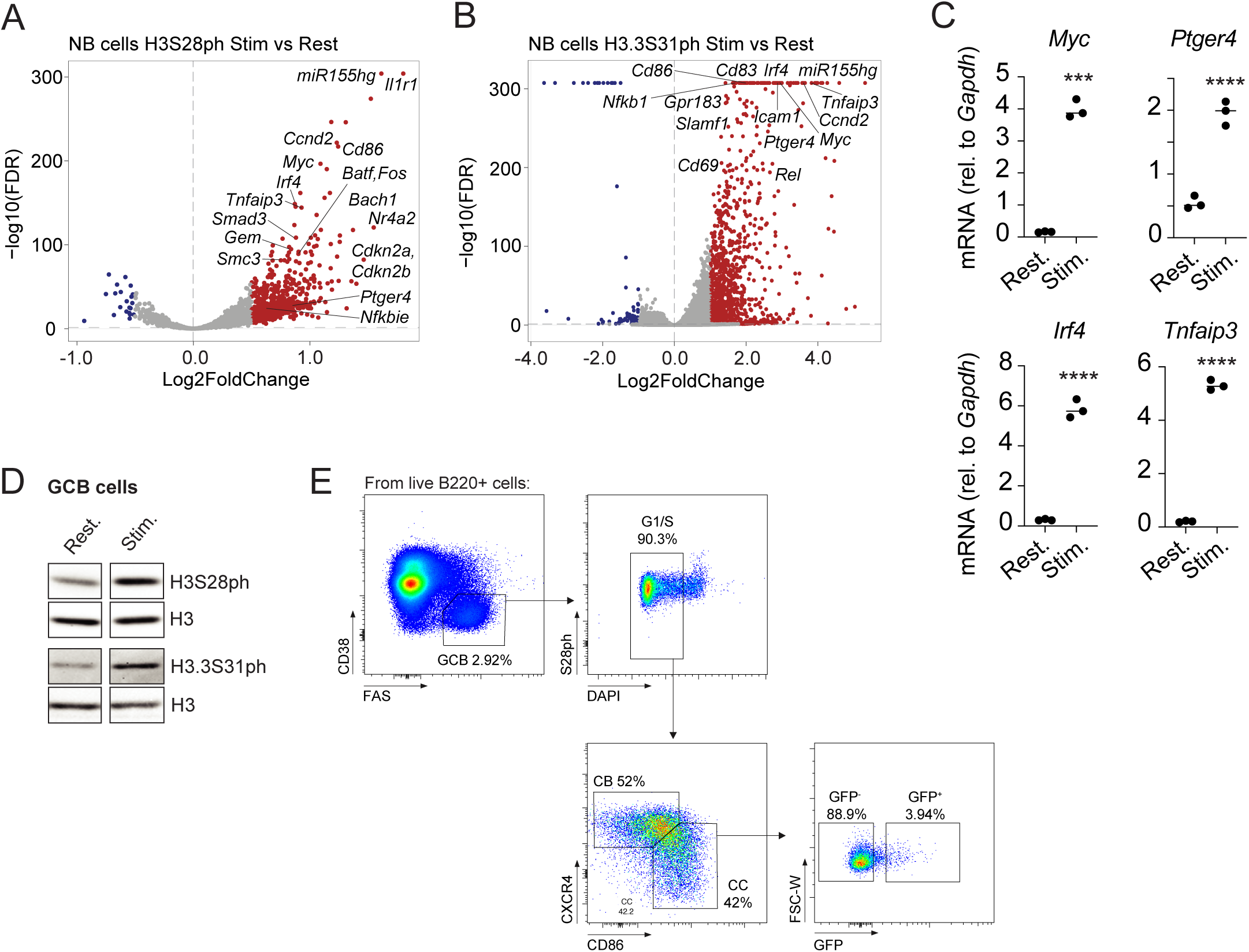
**(A, B)** Volcano plot of differentially phosphorylated targets between stimulated and resting naïve B cells (relevant B cell activation genes are labeled). (*n*=4). **(C)** Quantitative PCR of histone phosphorylation targets in resting and stimulated (conditions as in B) naïve B cells. (*n*=3, mean ± s.d. Unpaired *t* test (two-tailed), **** p-val <0.001, **** p-val <0.0001). **(D)** Western blot on resting and *ex vivo* stimulated (one hour anti-IgM, anti-IgG, anti-CD40 and IL-21) germinal center B cells (GCB), representative of 2 independent experiments. **(E)** Example gating strategy for the assessment of histone phosphorylation status on G1/S cells using the *Myc^GFP/GFP^* reporter mouse.

**Figure S3.**
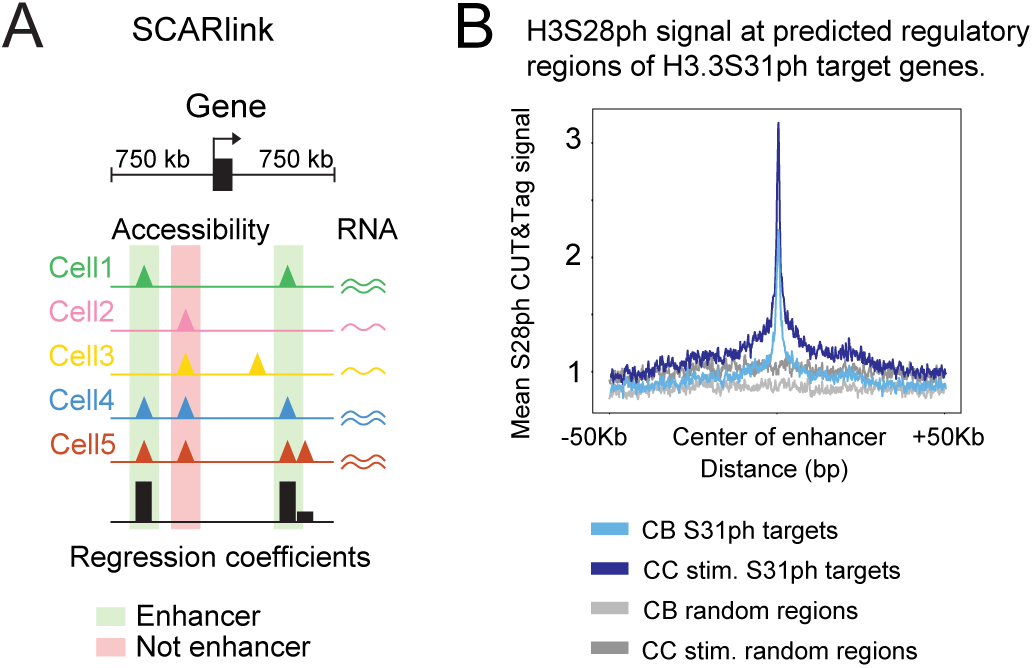
**(A)** SCARlink schematic. **(B)** Average H3S28ph CUT&Tag signal over SCARlink-identified regulatory regions of H3.3S31ph target genes and randomly selected regions.

**Figure S4.**
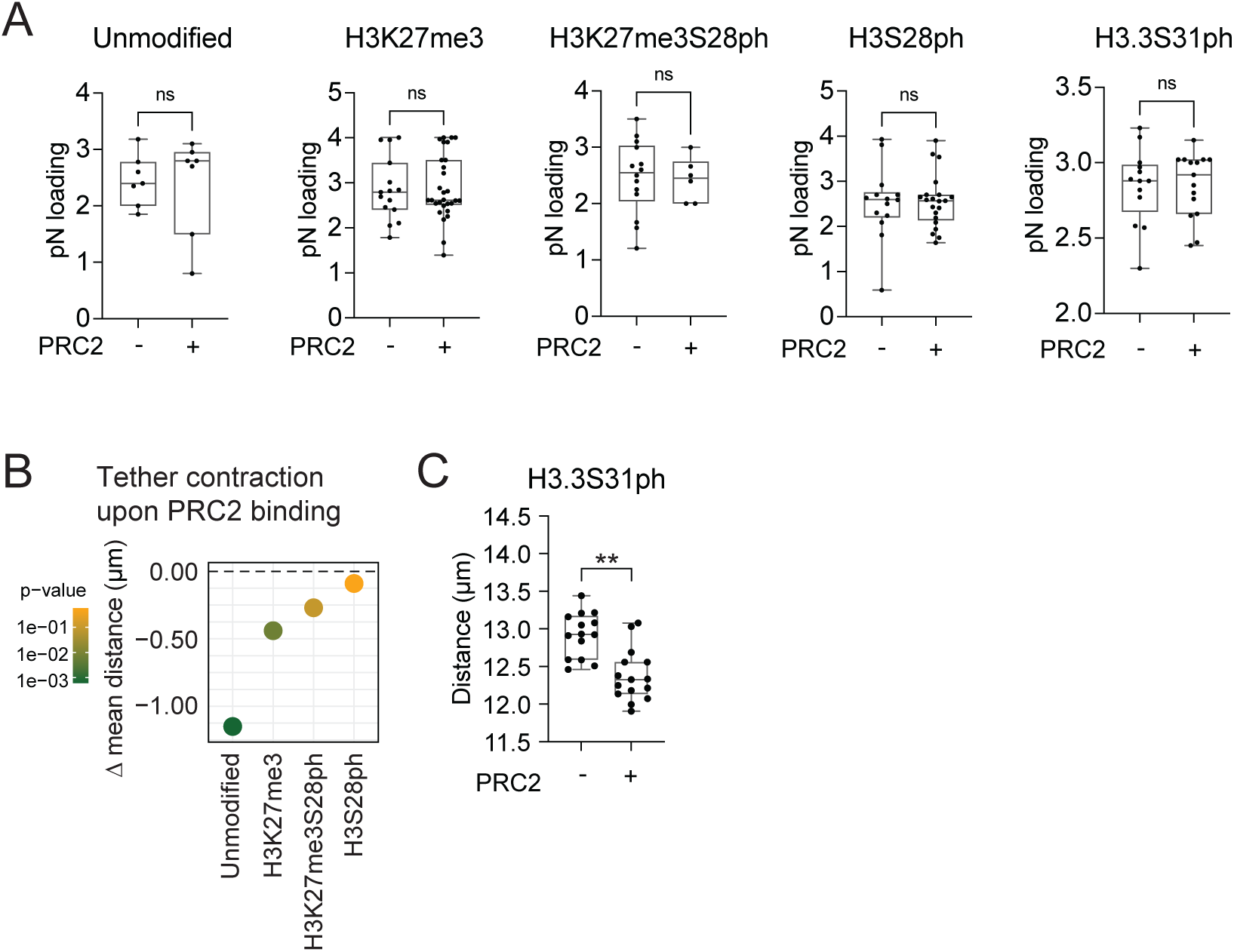
**(A)** Loading levels of the indicated designer octamer species onto single molecules of lambda DNA using force as a proxy. Each dot represents an independent DNA tether (Unmodified *n*=7, unmodified + PRC2 *n*=7, H3K27me3 *n*=15, H3K27me3 + PRC2 *n*=29, H3S28ph *n*=14, H3S28ph + PRC2 *n*=22, H3K27me3S28ph *n*=15, H3K27me3S28ph + PRC2 *n*=6, H3.3S31ph *n*=14, H3.3S31ph + PRC2 *n*=15. Unpaired Mann-Whitney *U* test (two-tailed). **(B)** Comparison of mean tether contraction upon PRC2 binding from measurements in Figure 4I. **(C)** Comparison of H3.3S31ph chromatin tether extensions at 3pN of force in the absence or presence of PRC2+AEBP2 complex (each dot represents a single chromatin molecule, H3.3S31ph *n*=14, H3.3S31ph + PRC2 *n*=15. Unpaired Mann-Whitney *U* test (two-tailed), ** p-val <0.01).

**Figure S5.**
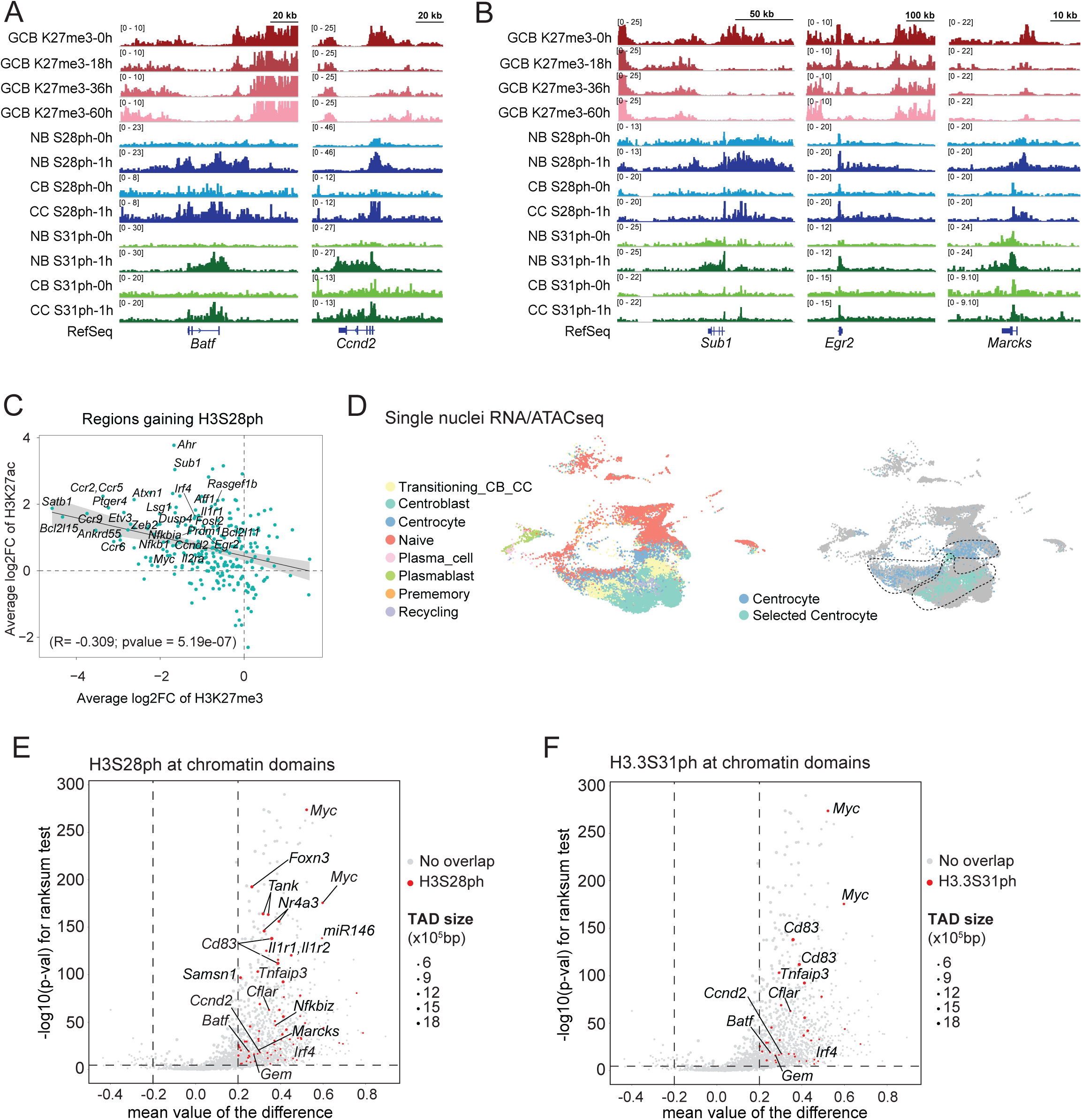
**(A, B)** ChIPseq and CUT&Tag tracks of naïve B cells (NB), centrocytes (CC), centroblasts (CB) and germinal center B cells (GCB), resting and stimulated for the indicated time points. **(C)** Plot depicting average log2 fold change of H3K27ac and H3K27me3 peaks overlapping peaks of increased H3S28ph after stimulation. **(D)** UMAP of 10x Multiome snATAC/RNAseq on germinal center B cells enriched with selected Myc^GFP+^ cells. Cell type annotations are based on expression profile datasets. **(E-F)** Differential predicted topologically associating domain (TAD) mean interactivity between selected centrocytes and centrocytes. Highlighted TADs overlap with H3S28ph (C) and H3.3S31ph (D) target genes.

**Figure S6.**
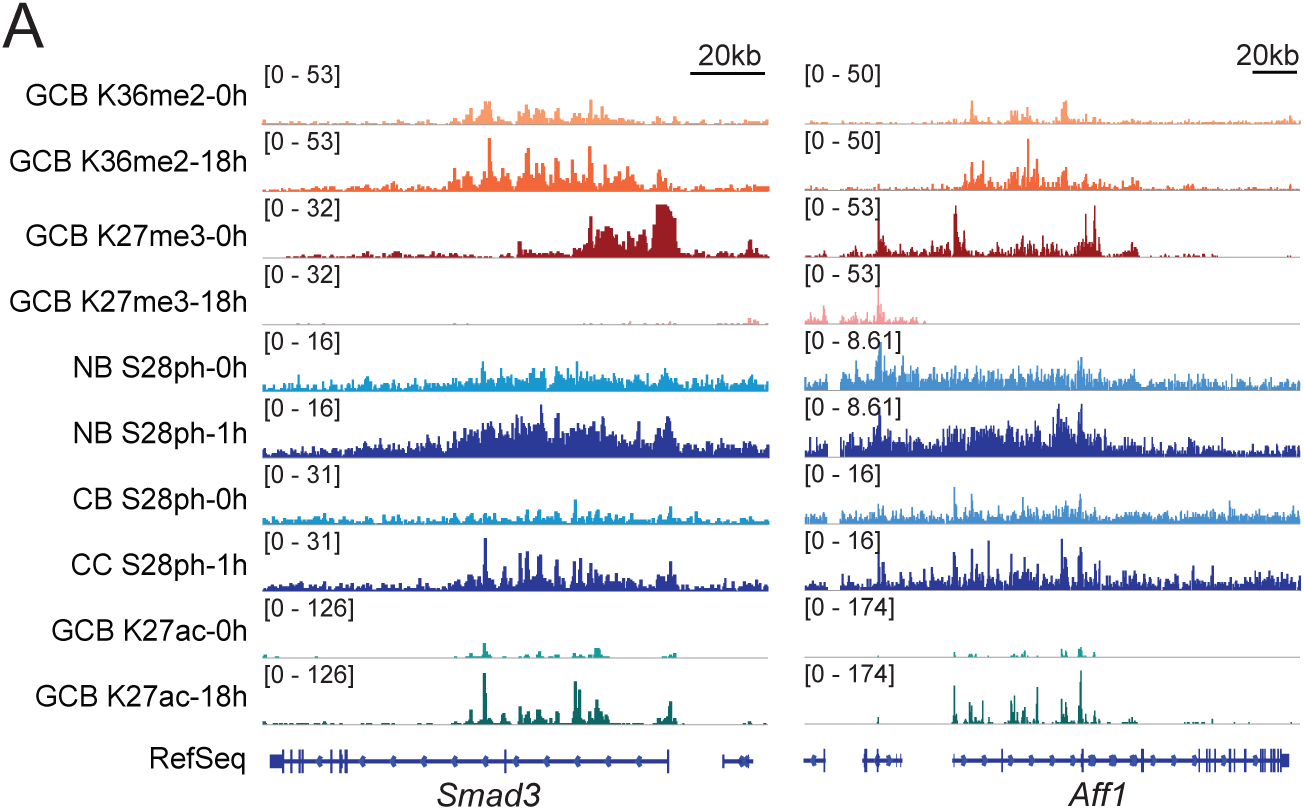
**(A)** Example IGV tracks for the indicated histone posttranslational modifications and resting and stimulated B cell populations.

